# A mid-level organization of the ventral stream

**DOI:** 10.1101/213934

**Authors:** Bria Long, Chen-Ping Yu, Talia Konkle

**Affiliations:** Department of Psychology, Harvard University; Department of Psychology, Stanford University; Phiar Technologies, Inc.

**Author notes:** Correspondence: Bria Long, Department of Psychology, Stanford University Jordan Hall, 450 Serra Mall Stanford, CA 94305.

## Abstract

Human object-selective cortex shows a large-scale organization characterized by the high-level properties of both animacy and object-size. To what extent are these neural responses explained by primitive perceptual features that distinguish animals from objects and big objects from small objects? To address this question, we used a texture synthesis algorithm to create a novel class of stimuli—*texforms*—which preserve some mid-level texture and form information from objects while rendering them unrecognizable. We found that unrecognizable texforms were sufficient to elicit the large-scale organizations of object-selective cortex along the entire ventral pathway. Further, the structure in the neural patterns elicited by texforms was well predicted by curvature features and by intermediate layers of a deep convolutional neural network, supporting the mid-level nature of the representations. These results provide clear evidence that a substantial portion of ventral stream organization can be accounted for by coarse texture and form information, without requiring explicit recognition of intact objects.

**SIGNIFICANCE STATEMENT:** While neural responses to object categories are remarkably systematic across human visual cortex, the nature of these responses been hotly debated for the past 20 years. In this paper, a new class of stimuli (“texforms”) is used to examine how mid-level features contribute to the large-scale organization of the ventral visual stream. Despite their relatively primitive visual appearance, these unrecognizable texforms elicited the entire large-scale organizations of the ventral stream by animacy and object size. This work demonstrates that much of ventral stream organization can be explained by relatively primitive mid-level features, without requiring explicit recognition of the objects themselves.

## INTRODUCTION

The ventral visual stream transforms retinal input into representations that help us recognize the categories of objects in the visual world (DiCarlo & Cox, 2007; Mishkin, Ungerleider, & Macko, 1983). For a few specific categories—faces, bodies, scenes, and visual words—there is a mosaic of highly-selective neural regions in the occipito-temporal cortex (Kanwisher et al., 1997; Downing et al., 2001; Epstein and Kanwisher, 1998; Cohen et al., 2000). However, other basic-level category distinctions (e.g., shoes vs. keys) can also be decoded from multi-voxel patterns in this same cortex (Haxby, 2001; Julian, Ryan, & Epstein, 2016). Even broader categorical distinctions reflecting the animacy and real-world size of objects are evident in large-scale spatial structure of occipito-temporal cortex (Chao et al. 1999; Konkle & Caramazza, 2013; Konkle & Oliva, 2012; Martin et al., 2007; for review see Grill-Spector & Weiner, 2014).

However, understanding the nature of the visual feature tuning underlying these ubiquitous categorical responses and their spatial organization across the cortex has proven notoriously difficult.

One key challenge is methodological: any measured neural response to recognizable object categories may actually reflect the processing of low-level image statistics, mid-level perceptual features, holistic category features, or even semantic associations that are not visual whatsoever (or some combination of these features). In other words, there is a continuum of possible representational levels that could account for neural responses to object categories. Within a classic view of the ventral visual hierarchy (Goodale & Milner, 1982; Kravitz et al., 2013; DiCarlo & Cox, 2007), there is broad agreement that low-level features are processed in early visual regions, and high-level, categorical inferences take place in later, downstream regions, including the anterior temporal lobe (e.g., Lambon Ralph et al., 2016). But, for the neural representations in intermediate occipito-temporal cortex, there is active debate about just how “high” or “low” the nature of the representation is.

On one extreme, some evidence suggests that the categorical neural responses are quite high-level, reflecting the *interpretation* of objects as belonging to a given category, rather than anything about their visual appearance per se (see Peelen & Downing, 2017 for a recent review). For example, when ambiguous moving shapes are identified as ‘animate’, they activate a cortical region that prefers animals (Castelli et al., 2003; Wheatley, Milleville, & Martin, 2007). Within the inanimate domain, hands-on training to treat novel objects as tools increases neural responses to these novel objects in tool-selective areas (Weisberg et al., 2006). In addition, differences between object categories persist when attempting to make them look as similar as possible (e.g., a snake versus a rope; Bracci & Op de Beeck, 2016; Kaiser et al., 2016; Proklova et al., 2016; but see Murty & Pramod, 2016 for critiques to this approach). These findings, and others from congenitally blind participants (e.g., He et al., 2013; Peelen et al., 2014; Striem-Amit & Amedi, 2014; van de Hurk, Van Baelen, & de Beeck, 2017; for a review, see Bi, Wang, & Caramazza, 2016) have led to the strong claim that visual features are insufficient to account for categorical responses in visual cortex (Peelen & Downing, 2017).

At the same time, a growing body of work demonstrates that neural responses in occipito-temporal cortex also reflect very low-level visual information. Retinotopic maps are now known to extend throughout high-level visual cortex (Brewer et al., 2005; Golomb & Kanwisher, 2012; Hasson et al., 2002; Larsson & Heeger, 2006; Levy et al., 2001; Rajimehr et al., 2014).

Furthermore, low-level visual features, like luminance and the presence of rectilinear edges, account for a surprising amount of variance in neural responses to objects (Baldassi et al., 2013; Nasr et al., 2014). More recently, some evidence suggests that recognizable objects and unrecognizable, locally-phase scrambled versions of objects yield similar neural patterns across occipito-temporal cortex (Coggan et al., 2016; but see Ritchie et al., 2017). Taken together, these results have led to an alternative proposal, in which the categorical responses of occipito-temporal cortex are solely a byproduct of simple visual feature maps and are not related to the categories per se (Andrews et al., 2010; see also Op de Beeck et al., 2008).

These two current viewpoints represent two prominent models of how to characterize the representations in occipito-temporal cortex. On an intermediate account, of course, neural responses in occipito-temporal cortex reflect tuning to visual features of intermediate complexity (e.g. Lerner et al., 2004; Tanaka et al., 2003; for reviews, see Lehky & Tanaka, 2016; Grill-Spector & Weiner, 2014). That is, it is mid-level features, combinations of which reflect the ‘shape of things’ (Haxby, 2001) that underlie categorical responses. However, neural evidence for a mid-level feature representation is sparse, in part because there is no widely accepted model of mid-level features. For example, is the basis set of this feature space derived from generic building blocks (i.e. Biederman, 1987) or from features tightly-linked to categorical distinctions (e.g., the presence of eyes)? As such, isolating mid-level representations and mapping their relationship to categorical responses is both methodologically and theoretically challenging.

Here, we approached this challenge by leveraging a new class of stimuli—“texforms.” Specifically, we used a texture synthesis algorithm to generate stimuli which preserve some texture and coarse form features from the original images, but lack outer contours and recognizable shape parts (e.g., handles, tails, eyes) (Freeman & Simoncelli, 2011; Long, Konkle, Cohen, & Alvarez, 2016; Long, Störmer, & Alvarez, 2017). Critically, people cannot identify what these stimuli are, looking to most people like “texturey-blobs” (see Figure 1). However, when prompted to guess if a texform is likely to be an animal or an object, big or small, participants are above chance (Long et al., 2016, 2017). Thus, this prior work provides behavioral evidence that there exist mid-level perceptual features that distinguish animals vs. inanimate objects and big objects vs. small objects. With this stimulus set, we are now poised to ask whether the features preserved in these texform stimuli are sufficient to drive neural differences between animals and objects of different sizes, and where along the ventral stream any differences manifest.

**Figure 1:**
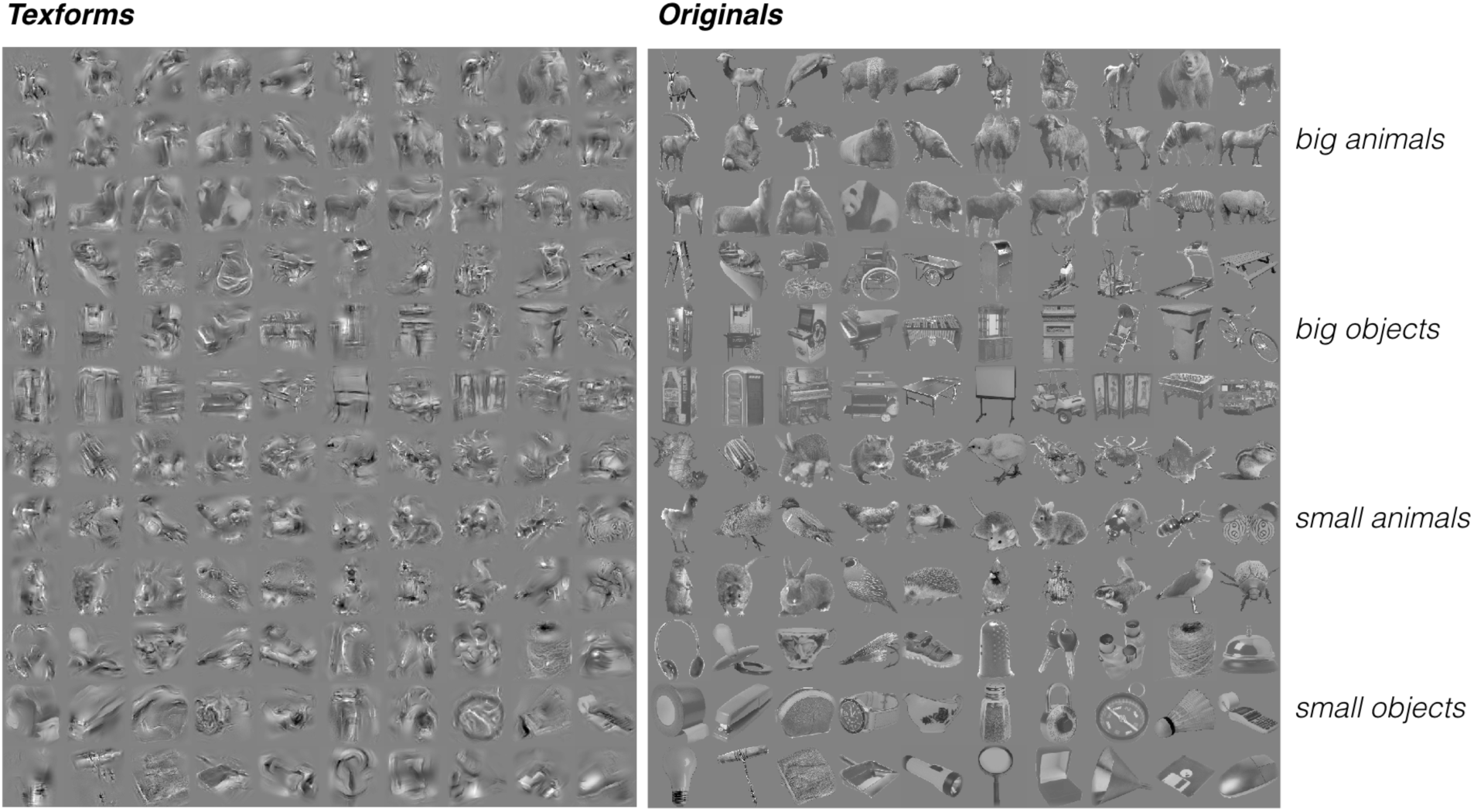
Stimuli: 120 texforms (right panel) were generated using a custom texture synthesis model (Freeman & Simoncelli, 2011) from recognizable pictures of 30 big objects, 30 small objects, 30 big animals, and 30 small animals (left panel). Stimuli were selected such that all texforms were unrecognizable at the basic level using online recognition experiments.

To anticipate our results, we find clear evidence that mid-level perceptual features are sufficient to drive the entire ventral stream organization by animacy and object size. Further, we compared how well a variety of models can predict the neural response patterns to these texform stimuli, finding that both intuitive curvature ratings and state-of-the-art deep convolutional neural net (CNN) unit tunings explain a large portion of the variance in neural pattern to both texforms and recognizable images. These results demonstrate that categorical responses in occipito-temporal cortex be explained to a large degree by relatively primitive mid-level perceptual features including texture and coarse form information, even driving purportedly “high-level” regions of the ventral stream. We propose that mid-level features meaningfully co-vary with high-level distinctions between object categories, and this relationship underlies the large-scale organization of the ventral stream.

## RESULTS

Observers viewed texform images of big animals, big objects, small animals, and small objects, followed by their recognizable counterpart images, while undergoing functional neuroimaging. **Figure 1** shows the full stimulus set. All images were presented in a standard blocked design, enabling us to examine the univariate effects of the two main dimensions (animacy and size) for both texforms and original image sets.

Additionally, we included a nested factor in the design related to texform classifiability. Specifically, for each texform, a classifiability score was calculated based on how well a separate set of observers could guess its animacy and real-world size. These scores were used to systematically vary how classifiable each block of texforms was (see Methods). Importantly, within this range of classifiability, there was no correlation with recognizability. In other words, a relatively well-classified image was no more likely to be recognized at the basic-level than a poorly-classified image. All original images were also presented in the same groups in yoked runs. This nested factor created a secondary, condition-rich design, enabling us to examine the structure of multi-voxel patterns to texforms and original images. Importantly, subjects in the neuroimaging experiment were never asked to classify the texforms (and were not even informed that they were viewing pictures of scrambled objects).

### Animacy and Object Size Topographies

To examine whether texforms elicited an animacy organization, we compared all animal and object texform univariate responses in each participant by plotting the difference in activation within a visually active cortex mask (all > rest, *t* > 2). Systematic response differences to animal versus object texforms were observed across the entire occipito-temporal cortex, with a large-scale organization in their spatial distribution. The same analysis was conducted using responses measured when observers viewed the original, recognizable images of animals and objects. The preference maps for both texforms and original animacy organizations are shown for a single subject in **Figure 2a**, and reveal a striking degree of similarity (see group topographies in **Figure S1**). Thus, even though there is an obvious perceptual difference between texforms and recognizable objects, they elicit matched topographies along the entire occipito-temporal cortex.

**Figure 2.**
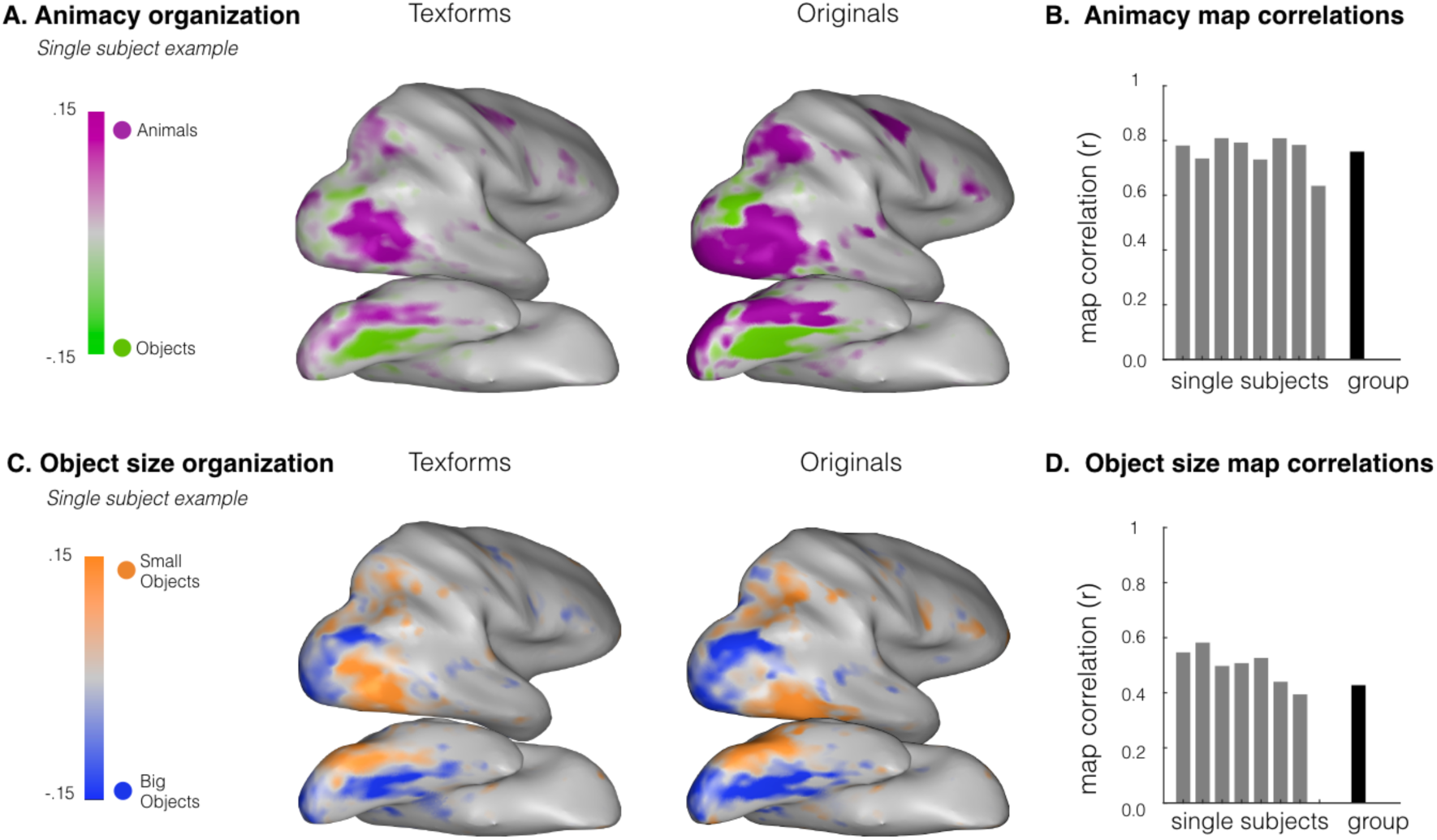
(A) Animacy topographies. The top row shows the animacy organization. Response preferences are plotted for animals versus objects for an example participant, considering texform images (left), and original images (right). (B) Animacy map correlations. The correlation between the original and texform response maps is plotted for all individual participants and averaged across all subjects for the animacy organization. (C) Object size topographies. Response preferences are shown for big objects versus small objects in an example participant, for texform images (left) and original images (right). (D) Object size map correlations. The correlation between the original and texform response maps is plotted for all individual participants and averaged across all subjects for the size organization.

To quantify this correspondence, we computed the correlation between the original and texform preference maps separately in each participant (Srihasam et al, 2014; see Methods). The map correlation coefficients for each participant and the average correlation coefficient for the group are plotted in **Figure 2b**. Overall, voxels in the occipito-temporal cortex had similar animacy preferences for recognizable and texform images in all subjects, resulting in a robust correlation at the group level (Average *r* = .76, *SD* = .06), that approached the noise ceiling (average noise ceiling across subjects, *r* = .83, paired t-test against noise ceiling, *t*(7) = 2.84, *p* = 0.025; see Methods).

Next, we used a similar analysis to examine whether texforms also elicited a real-world size organization. Given that the size organization is only found for inanimate objects, not animals (Konkle & Caramazza, 2013), we compared the responses to big objects versus small objects. Note that this yields half the power in the design to examine the object size organization relative to the animacy organization. Nonetheless, big and small object texforms elicited robust differential responses along the ventral stream, with a systematic large-scale spatial organization similar to that elicited by original images **(Figure 2c**; see group topographies in **Figure S1)**.

Quantitatively, moderate correlations between original and texform preference maps were found in all but one participant, resulting in map correlations at the group level that were not significantly different from the noise ceiling (**Figure 2d**; Average *r* = .43, *SD* = .21, Average noise ceiling across subjects *r* = .48; paired t-test against noise ceiling *t*(7) = −1.28, *p* = 0.24).

### Posterior-to-Anterior Analysis

Within a classic view of the ventral stream hierarchy, posterior representations reflect more primitive features while anterior regions reflect more sophisticated features. We next looked for evidence of this hierarchy with respect to the animacy and object size organizations, specifically examining whether original images evoked stronger animacy and size preferences than texforms in more anterior regions. To do so, we defined five increasingly anterior anatomical regions of occipito-temporal cortex (see Methods; **Figure 3A**).

**Figure 3.**
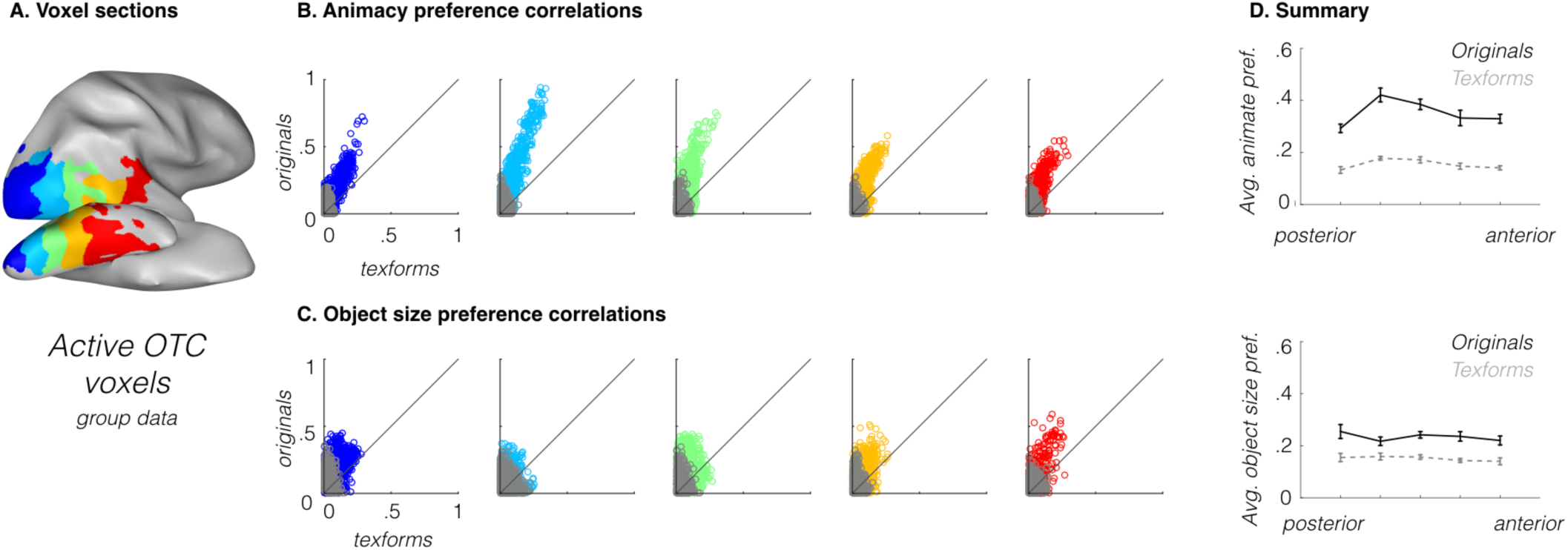
(A). Anatomical sections (shown here at the group level) from posterior to anterior in red to blue. (B) The animacy preferences elicited by texforms (x-axis) and by originals (y-axis) are plotted for each of the anatomical sections in the five subplots. Each point is a voxel. All points above the diagonal are voxels that show stronger animal/object preferences for original images than for texforms. Voxels where texforms and originals did not show the same preference are plotted in grey. (C) Object size preferences elicited by texforms (x-axis), and originals (y-axis) for each anatomical section, as in (B). (D). Average overall response preferences are plotted as a function of anatomical section for both originals (black solid line) and texforms (gray dashed line). Top panel reflects the animacy organization and summarizes (B); bottom panel reflects the object-size organization and summarizes (C). The averages were computed from data and anatomical sections defined at the individual subject level; error bars reflect between-subjects standard error of the mean.

We first looked at animacy preferences along this gradient to determine whether the category preference becomes increasingly larger for original images (vs. texforms) in more anterior regions. In **Figure 3b**, the five subplots compare the strength of animacy preferences elicited by originals vs. by texforms at the group-level. Each point is a voxel, and all points above the diagonal are voxels that show stronger animal/object preferences for original images than for texforms. These plots reveal stronger animate/object preferences for original images, compared to texforms, across all five anatomical sections (see Methods). This effect is quantified in **Figure 3d**, where the solid lines show the average preference strength across voxels for originals, and the dashed lines for texform images. If original images exhibited stronger category preferences in more anterior regions, this would be evident by an increasing difference between the solid and dashed lines. Instead, these two lines are relatively parallel (**Figure 3d** top panel; average beta difference for each section in animacy size preferences for original-texforms: *M*_S1_*=*.14, *M*_S2_ *=* .20, *M*_S3_= .18, *M*_S4_=.16, *M*_S5_ = .16; average rank correlation between these beta differences and anatomical sections, *r* = .06, t-test against zero, t(7)=0.31, *p* = 0.77). Thus, this analysis reveals that original images generate stronger category preferences than do texforms across *all* anatomical sections, and not only in more anterior ones.

When we conducted the same analyses on the object size distinction, we found the same pattern of effects. That is, original images elicited stronger big/small object preferences than texforms across all anatomical sections, and this difference was relatively consistent from posterior to anterior sections (**Figure 3E**, middle panel), average beta difference for each section in object size preferences for originals – texforms: *M*_S1_*=* .09, *M*_S2_ *=* .05, *M*_S3_= .08, *M*_S4_= .08, *M*_S5_ = .07; average rank correlation between these beta differences and anatomical sections, *r* = −.15, t-test against zero, *t*(7) = −0.59, *p* = 0.57). In other words, we found little if any evidence for the pattern of results that might be expected from a simple visual hierarchy, in which texforms and original neural responses matched in posterior areas, but diverged in anterior areas. Instead, the difference in animacy/size preferences to originals versus texforms remained relatively constant across the full posterior-to-anterior gradient.

One possible factor that might influence the interpretation of this result is about the overall neural activity: perhaps original images simply drive all voxels along the ventral stream more than texforms, and thus the greater animacy/size preferences we observe for original images actually reflects better signal-to-noise. On such an account, original and texform organizations may be even more similar to each other than we have measured. To examine this possibility, we analyzed the overall magnitude of the neural response to originals vs. texforms along this posterior to anterior axis, averaging across all animacy/size conditions. Overall, voxels were driven relatively similarly by both texforms and originals images across all anatomical sections, though if anything, recognizable images generated slightly more overall activity than texforms in the more *anterior* sections (average beta difference between originals and texforms in each section; *M*_S1_*=* −0.02, *M*_S2_ = .04, *M*_S3_ = .08, *M*_S4_ = .08, *M*_S5_ = .13, average rank correlation between beta differences and anatomical sections, *r* = .64, t-test against zero, *t*(7) = 4.51, *p* = 0.003). Thus, it was not the case that originals images elicited stronger overall responses everywhere, and response magnitude is unlikely to explain away the result that original images elicit stronger animacy/object and big/small preferences across the ventral stream. We consider several explanations for this difference in preference strength between texforms and originals in the General Discussion.

In sum, both texforms and recognizable images generated large-scale topographies by animacy/size throughout the entire ventral stream, with recognizable images generating overall stronger category preferences (without difference in overall response magnitude). We did not find strong evidence for a hierarchy of representations that differentiated between texforms and recognizable images. Instead, our results point towards mid-level features as a major explanatory factor for the spatial topography of object responses along the entire occipito-temporal cortex.

### Tolerance to Retinal Position

Given the extensive activation of these texforms along the ventral stream, one potential concern is that these texform topographies may reflect simple retinotopic biases that also extend throughout this cortex, rather than mid-level feature information per se. For example, if animal texforms happen to have more vertical information in the lower visual field, and object texforms have more horizontal information in the upper visual field, then such low-level differences might account for the responses observed in the main experiment. To test this possibility, we conducted a second experiment in which a new group of observers were shown the same stimuli (both texforms and recognizable images) but each image was presented separately above and below fixation. If animacy and size preferences are maintained over changes in visual field position, this provides evidence against a simple retinotopic account.

We used a conjunction analysis to isolate voxels that showed stable category preferences regardless of the retinotopic location of the images. To do so, animal vs. object preferences were computed when original images were presented in the upper visual field location and separately for the lower visual field location. We retained voxels that showed the same category preference (e.g., animals > objects or objects > animals) when stimuli were presented in the upper visual field *and* when stimuli were presented in the lower visual field. The percent of retained voxels, relative to the total set of visually-active occipito-temporal cortex voxels, was computed for both the animacy and object size distinctions, for both original and texform images, separately in each participant.

When subjects viewed the original images, we found that 77% (*SD* = 5%) of voxels in occipito-temporal cortex showed location-tolerant animacy preferences, and 56% (*SD* = 9%) of voxels showed location-tolerant object size preferences. When subjects viewed texforms, we found that 59% (*SD* = 15%) of occipito-temporal voxels showed location-tolerant animacy preferences, and 49% (*SD* = 7%) of voxels showed location-tolerant object size preferences.

Thus, both recognizable images and texforms elicited animacy and object size preferences that were largely tolerant to changes in visual field position.

Next, we assessed the similarity of category preferences elicited by texform and originals within location-tolerant voxels. To do so, we again conducted map correlations, restricting our analysis to voxels that showed consistent category preferences for original images. These conjunction topographies for a single subject are shown in **Figure 4a** and group topographies are shown in **Figure S2.** Texform and original topographies again showed a high degree of spatial correspondence, evident in single subjects and at the group level (Animacy: average *r* = .75, *SD*= .09, *t(*7) = 13.93, *p* < 0.001; Size: *r* =. 38, *SD* = .14, *t(*7) = 7.03, *p* < 0.001; see **Figure 4b**).

**Figure 4.**
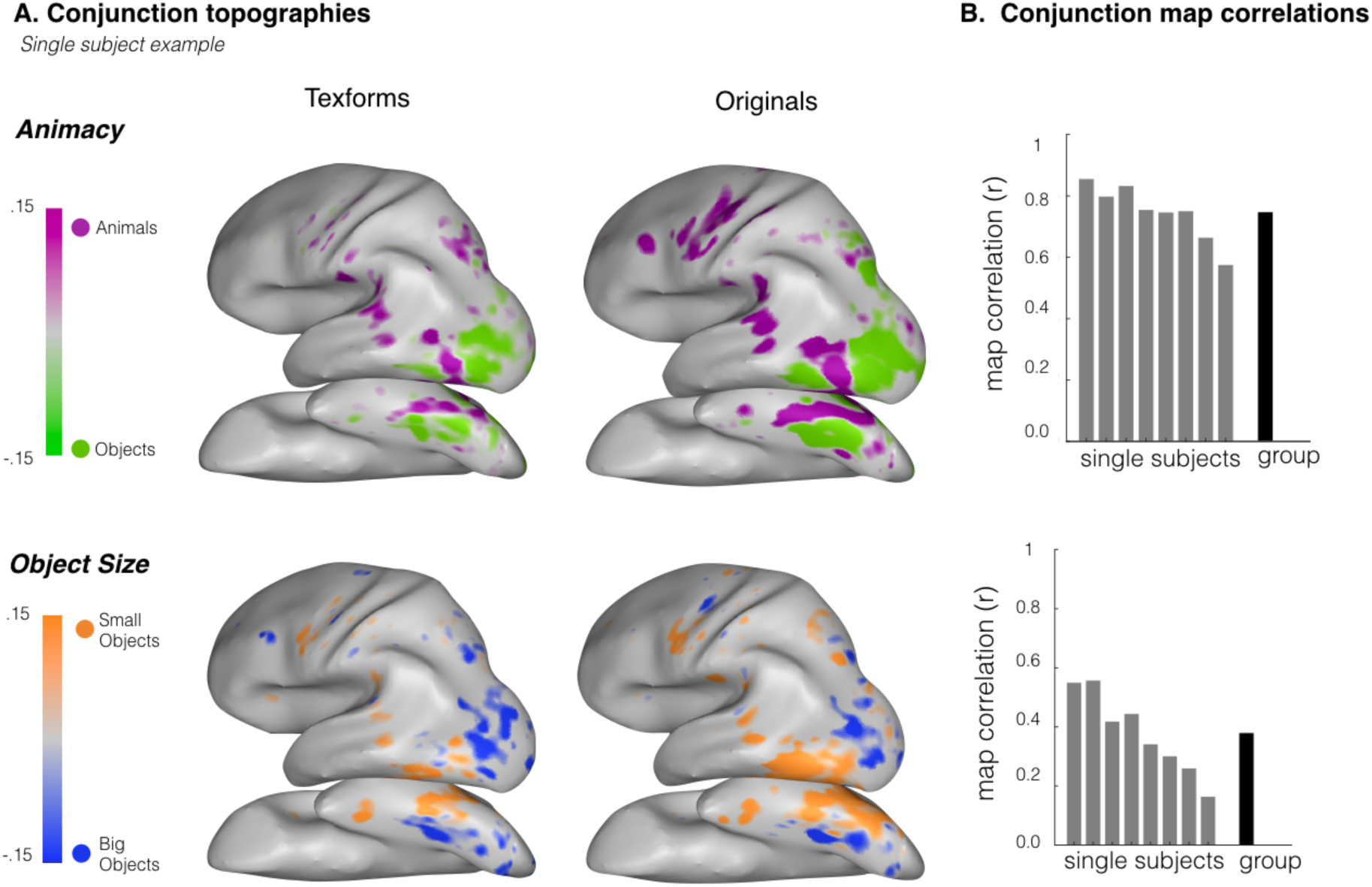
(A). Group conjunction topographies. Average category responses when texforms/originals (left, right) are presented in the upper and lower visual field. Topographies are restricted to voxels that show the same category preference regardless of the stimuli’s location in the visual field and are shown separately for animacy (upper panel) and size (lower panel). (B) Conjunction map correlation values (y-axis) are plotted for each individual subject (x-axis) and at the group level separately for animacy (upper panel) and object size (lower panel) contrasts.

Compared to the initial experiment, the organizations found here are sparser, particularly for the object size distinction. This may indicate a stronger contribution of retinotopic contributions for the object size relative to the animacy distinction, or may simply reflect the lower signal-to-noise in the object size analysis (for which only half the data are used). Nonetheless, these results demonstrate that these topographies reflect mid-level information that is tolerant to changes in visual field position, replicating and extending the primary finding.

### Predictive Modeling: Texforms

We next aimed to provide insight into the nature of the mid-level features that actually drive these animacy and size responses. To do so, we compared how well a variety of models predict the multi-voxels pattern similarity across occipito-temporal cortex (Jozwik et al., 2016; Nili et al., 2014; Khaligh-Razavi & Kriegeskorte, 2014).

To do so, we first constructed representational dissimilarity matrices (RDMs) in occipito-temporal cortex using data from the richer condition structure nested in our experiment design. Recall that every time observers saw a block of texform images, this block was comprised of a set of texforms from one of six levels of classifiability. The more classifiable the texform, the better a separate group of norming participants were able to guess that this texform was generated from an animal vs. an object, or from a big vs. small thing in the world (see Methods; **Figure S3**). Examples of well-classified and poorly-classified texforms (and their accompanying original counterparts) are shown in **Figure 5a (**see **Figure S3** for data from behavioral judgements). The fact that participants are above chance at these animacy/size classification tasks is informative: the texture-synthesis algorithm does not preserve enough visual features for participants to identify objects at a basic level, but retains sufficient information for observers to make guesses about their animacy and size in the real-world (Long et al., 2016; 2017).

**Figure 5.**
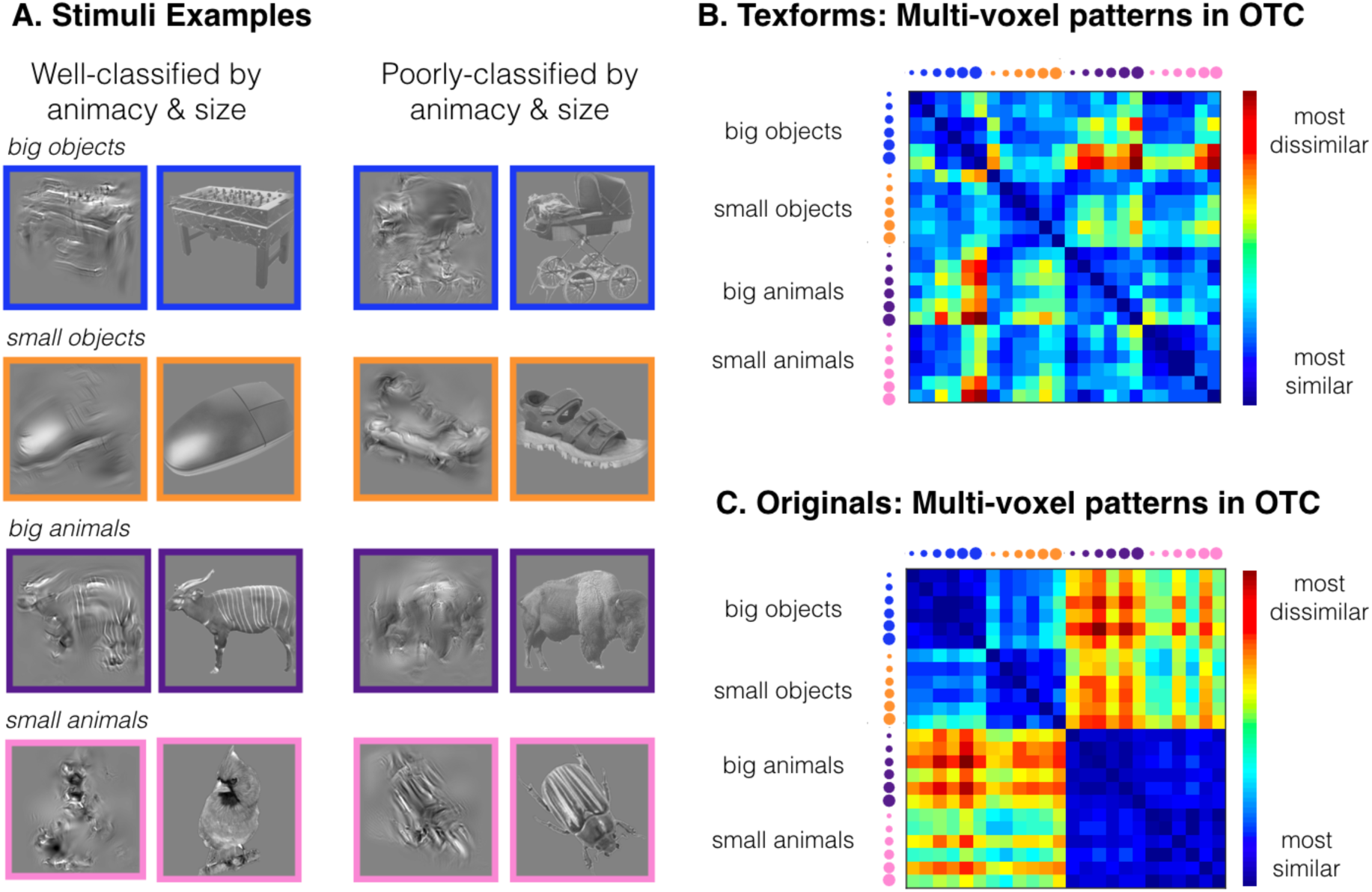
(A). Examples of texforms (and their corresponding originals). Stimuli on the left are examples from the best classified group of texforms, while stimuli on the right are examples form the worst classified group of texforms. (B). Representational dissimilarity matrix obtained from neural patterns in active occipito-temporal cortex for texforms and (C) their corresponding originals. The size of the dots corresponds to the degree to which the texforms in this nested group of images were classifiable by their animacy and real-world size. Data are scaled such that, in both cases, the most dissimilar values are red and the least dissimilar values are blue.

**Figure 5b** shows the similarity in the neural patterns elicited by texforms, and the corresponding patterns for originals are shown in **Figure 5c.** The texform RDM has some gradation in terms of the levels of texforms’ classifiability, which by inspection shows that more classifiable texforms are more dissimilar from each other. By comparison, the structure in responses to recognizable images is more categorical in nature, with a clear animate/inanimate division that is visually evident in the quadrant-structure of the RDM and with a weaker but visible big/small object division in the upper left quadrant.

What features best predict this neural similarity structure generated by texforms? We tested a variety of models following the procedure introduced by Jozwik et al., 2016. The key outcome measure was the degree to which a cross-validated feature model predicted the observed neural data in each subject (using rank correlation Kendall Tau-α, T_A_). All model performance is put in the context of the neural noise ceiling, reflecting how well a given subject’s RDM can predict the group RDM (see Methods; Nili et al., 2014), and shown in **Figure 6**.

**Figure 6.**
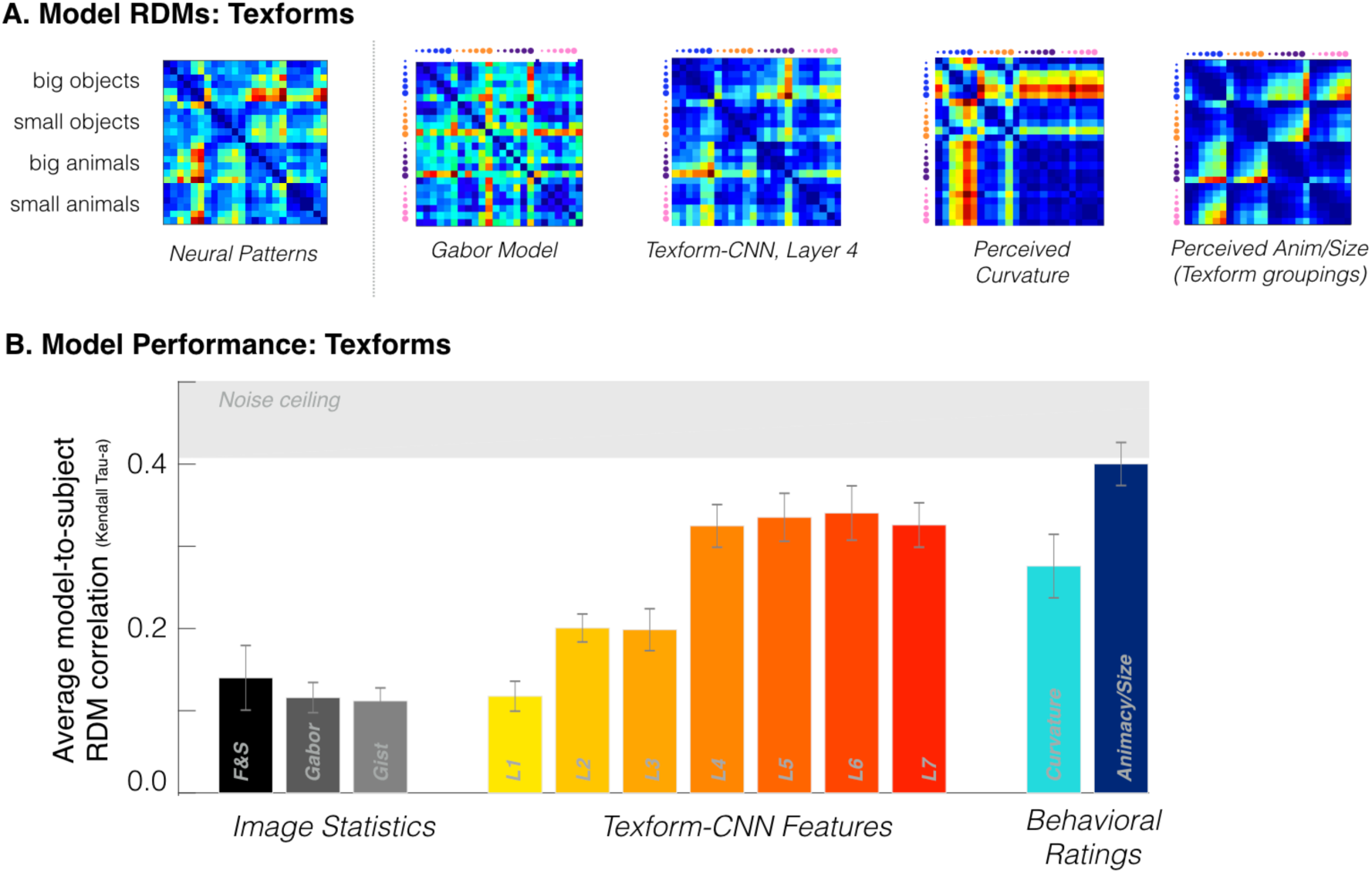
(A) Neural patterns in response to texforms (same as in Figure 4B) and predicted neural dissimilarities for selected models obtained through the same cross-validation procedure. (B) Average predicted model correlation (kendall tau-a) with individual subjects’ neural patterns in OTC. The bars show different models, from left to right: Freeman and Simoncelli texture model (black), gabor model (dark gray), gist model (light gray), AlexNet features layer 1 through layer 7 (yellow to red), curvature behavioral ratings (light blue), and animacy/size behavioral ratings (dark blue). Data is plotted with respect to the noise ceiling, shown in light gray.

#### Image Statistics

First, we examined how well combinations of low-level image statistics could account for the observed neural structure. Can combinations of texture synthesis parameters computed in local pooling regions explain the structure in the neural patterns? We found that weighted linear combinations of the texture-synthesis model parameters predicted relatively little variance in the neural patterns (T_A_ = .14). Consistent with this result, other models based on low-level image statistics also only captured a small amount of variance (Gabor model, T_A_ = .12; Gist model, T_A_ = .11; Oliva & Torralba, 2006). Thus, linear combinations of relatively simple visual features were not sufficient to predict the neural responses to texforms, despite the fact that these image features were used to generate the texforms. This implies that the mid-level features that elicit this multi-voxel pattern structure are not explicit in the parameters of the texture synthesis model or their simple linear combinations.

#### Convolutional Neural Nets (CNNs)

We next tested models constructed from deep CNN unit responses, reflecting the state of the art in predicting neural responses to objects (e.g., Güclü & van Gerven, 2015; Khaligh-Razavi & Kriegeskorte, 2014). To do so, we extracted the feature representations throughout all layers of a CNN (Krizhevsky et al., 2012) in response to texforms. Note that while this CNN was pre-trained to categorize one thousand object categories, it was not specifically trained (or tuned) on any of the texform images or their recognizable counterparts.

Models constructed from representations in the earliest layers of a CNN performed poorly, similar to the models based solely on image statistics. However, predictive ability increased through the first few convolutional layers, plateauing around convolutional layers 4 and 5 (Layer 4, T_A_ =.32; Layer 5, T_A_ =.34). Thus, the variation in neural patterns to different groups of texforms was relatively well captured by responses in mid-level convolutional layers of a deep CNN. These results reveal that mid-level features captured by these intermediate CNN layers can explain the variation in neural patterns to different groups of texforms.

#### Curvature Ratings

We next asked how well perceived curvature ratings could explain this neural structure, based on behavioral evidence that boxy/curvy ratings distinguish between animals, small objects, and big objects (Long et al., 2016; 2017; Long & Konkle, 2017), and in line with a growing body of work implicating curvature as a critical mid-level feature in ventral stream responses (e.g. Nasr et al., 2014; Ponce et al., 2017; Srihasam et al., 2014; see **Figure S4** for behavioral judgments). We found that this simple, one-dimensional model based on curvy-boxy judgments was able to predict the structure moderately well (T_A_ =.28), capturing almost 50% of the variance in the neural patterns elicited by texforms.

#### Animacy/Size Classification

As a sanity check, we examined the performance of a behavioral model constructed directly from the classification scores used to group the texforms into the nested conditions by classifiability. We expected this model to perform well, as we built this structure into our experiment design. Overall, we found that these animacy-size judgments were able to predict the structure of texform responses near the noise ceiling (average subject RDM-to-model correlation, T_A_ =.40; noise ceiling T_A_ = [.40—.50]). This result confirms that the neural patterns in response to texforms varied as a function of the classifiability of the texforms; groups of texforms that were better classified by their animacy/size elicited more distinct neural patterns.

#### Modeling in Early Visual Cortex (EVC)

Finally, given that we generated texforms using a texture synthesis model that explains variance in V2/V4 (Freeman et al., 2013; Okazawa et al., 2015), we explored which feature spaces explained variance in in early visual cortex by applying the same analytic method (see **Figure S5**). Consistent with prior work, we found that Gabor-based models explained the most variance in early visual cortex, whereas models based on perceptual properties (e.g., perceived curvature) or category-based models explained less variance. Thus, these results suggest that texforms elicit neural patterns in occipito-temporal cortex that are different from those elected in early visual cortex.

A complete summary of these modeling results is shown in **Figure 6**. To visually inspect the structure captured by the different models, **Figure 6a** shows the predicted neural RDM from the best-performing models. The overall performance for all models, reflecting the average model-to-subject RDM correlation, is shown in Figure **6b**. Taken together, these analyses show that models based on intuitive curvature ratings and intermediate layers of a deep CNN captured this neural structure relatively well, while models based on early CNN layers and simple image statistics were insufficient. Broadly, these modeling results provide computational support for the mid-level nature of this neural representation, and help triangulate the kinds of features that drive neural responses to texforms in occipito-temporal cortex (i.e. curvy/boxy mid-level features of intermediate complexity).

### Predictive Modeling: Recognizable Images and Cross-Decoding

For completeness, we also compared how well the same set of models could predict the structure of neural responses to original images, with the results summarized in **Figure 7a and 7b.** Overall, we found a similar pattern of results as we did with texforms. That is, models based on simple image statistics performed only moderately well, failing to capture most of the variance (Freeman & Simoncelli features, T_A_ =.13; Gabor features, T_A_ = .14; Gist model, T_A_ = .14; Neural noise ceiling [T_A_ = .74—.78]). However, the feature representations elicited in deep CNNs by recognizable images almost fully predicted these neural patterns. Model performance again reached a plateau when features came from intermediate layers in response to recognizable images (Layer 4, T_A_ = .65; Layer 5, T_A_ = .65). Interestingly, as with texforms, a model based on curvy-boxy judgments of the original images also accounted for a substantial portion of this variance (T_A_ = .43). Finally, a categorical model derived from the actual animacy and size of the images performed moderately well, though it was relatively far from the noise ceiling (T_A_ = .42), consistent with prior work pointing out the more graded similarity structure present in occipito-temporal cortex (Khaligh-Razavi & Krigeskorte, 2014).

**Figure 7.**
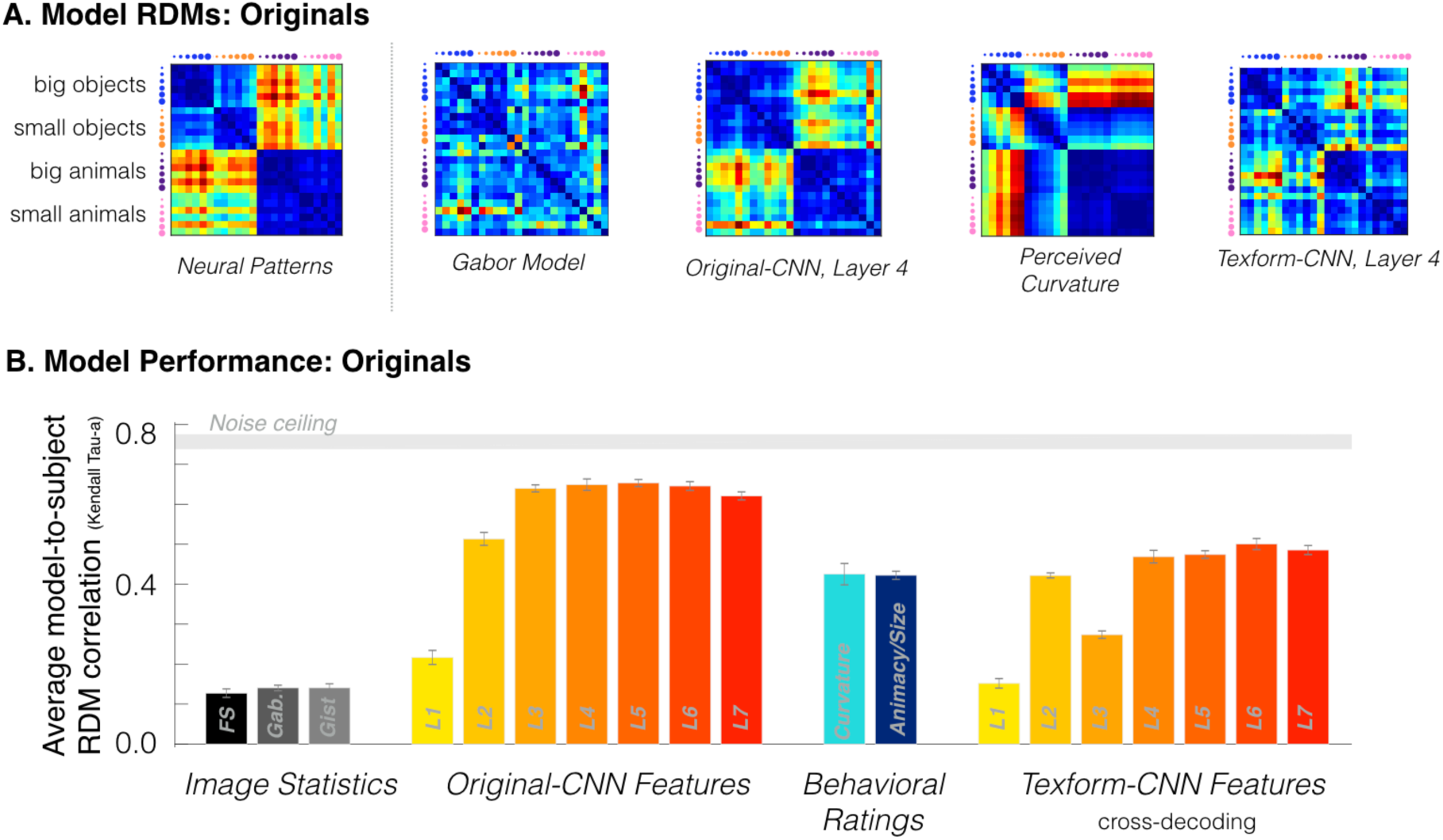
(A) Neural patterns in response to recognizable images (same as in Figure 4C) and predicted neural dissimilarities for four models obtained through the same leave-one-out cross-validation procedure. (B) Average predicted model correlation (kendall tau-a) with individual subjects’ neural patterns in OTC. The bars show different models, from left to right: Freeman and Simoncelli texture model (black), gabor model (dark gray), gist model (light gray), AlexNet features layer 1 through layer 7 extracted from the original images (yellow to red), curvature behavioral ratings (light blue), and animacy/size behavioral ratings (dark blue), and AlexNet features layer 1 through layer 7 extracted from the texform images (yellow to red). Data is plotted with respect to the noise ceiling, shown in light gray.

Taken together, the predictive power of the intermediate CNN layers and the curvature ratings also suggest that mid-level representation underlie a substantial component of neural responses to recognizable images. We next performed a stronger test of this argument by conducting a cross-decoding analysis. Specifically, we examined whether neural responses to original images could be predicted using the CNN features extracted from texforms. In other words, we tested whether the neural similarity of original images could be predicted from deep neural network responses to the texform counterparts of each original image. Indeed, texform-CNN-features were able to predict much of the RDM structure elicited by recognizable images (Layer 4, T_A_ = .47, **Figure 7b**). This cross-decoding analysis further supports the idea that the neural responses to recognizable objects are driven substantially by the mid-level feature information instead of by ‘high-level’ features, such as recognizable object parts or identity information. We consider several possible interpretations of this result in the discussion.

## GENERAL DISCUSSION

We employed a novel stimulus set—texforms—to examine if and how mid-level features contribute to the large-scale organization of the ventral stream. We found that (1) texform stimuli were sufficient to elicit animacy and size topographies along occipito-temporal cortex, well into what are classically considered more high-level object-selective areas, (2) these mid-level topographies were not inherited from low-level retinotopic biases, as they generalized over visual field position, (3) the similarity structure of the neural representations to both texforms and recognizable images was best predicted by intermediate layers of a deep CNN, with a simple curvy-boxy perceptual axis explaining a modest amount of the structure, and (4) texform model features were able to account for a substantial amount of the neural similarity structure elicited by original recognizable images.

Taken together, these findings establish that mid-level feature differences can drive extensive categorical neural responses along the ventral stream and underlie the topographic organization by animacy and object size. Broadly, these results inform the debate about the nature of object representation in occipito-temporal cortex: First, they challenge a simple conception of the visual hierarchy, as relatively primitive texforms drove category differences in what is typically considered “high-level” visual cortex. Second, they highlight that curvature co-varies with broad category distinctions and provide an intuitive description of the kind of mid-level featural information represented in this cortex. Below, we discuss the implications of these findings for models of the ventral stream, the role of curvature in ventral stream organization, and the features that could account for the gap in neural responses to texforms vs. originals.

### Implications for models of the ventral stream

There are two main observations to note about the texform topographies, each with separate implications for the nature of representation along the ventral stream. The first observation is that the neural differences between different kinds of texforms are detectable at all. Consider Figure 1—these stimuli all look like texturey-blobs. Participants have no idea what they are seeing, or even that there are different kinds of things here. One real possibility was that the differences between the texforms would be far too subtle to drive any measurable differences in brain responses, especially measured with fMRI. However, the data show that the visual system is not only tracking these incoming textures, but it is also triggering specific neural responses that cleanly align with the animacy and the object size organizations. These data provide strong evidence that these regions do not require specific tuning to recognizable animal parts, for example, like eyes and tails, or recognizable object parts like handles, or even clear bounded contours at all, in order to trigger these neural response differences between animals and objects of different sizes. Instead, these data point to evidence that a more statistical and primitive level of features support broad category distinctions along the ventral stream.

The second observation is that these texform topographies actually extend much farther anterior than one might expect on a classic view of the ventral stream as a hierarchy. On such an account, visual areas exhibit selectivity to increasingly complex visual features (Felleman & Van Essen, 1991; Rust & Dicarlo, 2010; for a review, see DiCarlo & Cox, 2007), where it is typically assumed that each transformation *discards* information from the previous stages. For example, specificity along the shape dimension is thought to come at the expense of sensitivity to other dimensions (e.g. position). However, it is increasingly clear that the visual transformations along the ventral stream may not be as ‘lossy’ as initially theorized (e.g. Hong et al., 2016; Schwarzlose et al., 2008). Consistent with this emerging view, we find that mid-level features drive categorical responses throughout the entire ventral stream, and are not limited to posterior areas previously implicated in processing curvature (e.g. Kourtzi & Connor, 2011). These empirical findings support a growing view in which higher-level visual cortex is sensitive to features that span multiple levels of representation (e.g., Grill-Spector & Weiner, 2014): anterior regions may retain their sensitivity to low and mid-level features while also becoming increasingly tolerant to complex stimulus transformations.

### The role of curvature in ventral stream organization

We found that the similarity structure of neural responses across occipito-temporal cortex was not only well predicted by intermediate and later layers of multi-million-parameter convolutional neural net models, but also by a single dimension of perceived curvature. This was true for both original images and unrecognizable texforms. What is special about curvature? We have previously speculated that big objects tend to be boxier because they must withstand gravity while small objects tend to be curvier as they are made to be hand-held, whereas animals have few if any hard corners and are the curviest (Konkle & Oliva, 2011, Levin et al., 2001; Long et al., 2017). Thus, there is an ecological (non-arbitrary) relationship between curvature and category. In recent work, we have found direct evidence for this link (Long et al., 2016; 2017; in press): the curviest texforms tend to be perceived as animate, and the boxiest texforms tend to be perceived as big, inanimate objects. Thus, the perceptual axis from boxy-to-curvy seems to meaningfully align with the broad category divisions between animals and objects of different sizes.

Based on these sets of results, we suggest that ventral stream responses are organized according to mid-level feature maps that meaningfully co-vary with high-level, categorical dimensions. That is, the level of representation in the neural populations is relatively visual/statistical in nature, but the large-scale topography is still reasonably described by high-level animacy and object size distinctions. That there are animacy and object/size differences doesn’t mean this whole cortex is solely involved in computing these high-level factors; rather, we suggest that mid-level shape space has major structure related to these two high-level dimensions.

Of course, one of the big unanswered questions about this relationship is the direction of causality: Are broad category distinctions like animacy and size learned on top of initial curvature biases in the visual system (Ponce et al., 2017)? Or, do mid-level curvature biases emerge as a byproduct of “top-down” influences or network-based constraints (Konkle & Caramazza, 2016), which enforce these particular broad category distinctions? Note that the present data cannot speak to the directionality of these mid and high-level factors, only to the existence of the link between them.

While the origins of curvature biases remain a source of debate, these results join a long history of research documenting the importance of curvature in ventral stream responses. An elegant series of studies have demonstrated the explanatory power of curvature in explaining single unit responses in V4 (Carlson et al., 2011; Pasupathy & Connor, 2001, 2002; Yau et al., 2012). Other work has shown systematic preferences for curvilinear versus rectilinear stimuli in different category-selective regions in infereo-temporal cortex (Caldara et al., 2006, 2009; Nasr, Echavarria, & Tootell, 2014; Rajimehr et al., 2011; Srihasam et al., 2014; but see Bryan, Julian, & Epstein, 2016). Most recently, curvature has been proposed as a proto-organizing dimension of the ventral visual stream (Srihasam et al., 2014; Ponce et al., 2017), and specific curvature-preferring patches have been discovered in macaques (Yue et al., 2014).

However, it is important to interpret this convergence of “curvature” findings with some caution. While curvature is an intuitive concept, it surprisingly hard to define computationally. As a result, curvature has been operationalized in relatively different ways across research groups. While we focused on a curvy-boxy dimension, others have defined the endpoints of this dimension as wavy-versus-straight (Srihasam et al., 2014; Ponce et al., 2017). Others have focused on the distinction between round-versus-rectilinear shapes (Yue et al. 2014). Thus, while curvature is clearly a productive property of shape space, it remains unclear if this collection of findings derives from a single, underlying construct of curvature.

### Differences between Texform and Original Responses

While we have emphasized the degree and extensiveness of the texform topographies, they are certainly distinguishable from the neural responses evoked by original, recognizable images. First, original images generated stronger categorical responses than texforms across the entire ventral stream in both the univariate effects and in their multi-voxel patterns. Second, CNN features extracted in response to *original* images were necessary to best predict the neural structure generated by recognizable images; texform-CNN-features did well but did not reach the same level as the original-CNN-features. These modeling results imply that the recognizable images contain additional visual features, captured by CNNs, which contribute to neural responses in occipito-temporal cortex. Below, we speculate about what visual features account for this “gap” between texform and original images, and how this relates to the debate about the nature of the representations along the ventral stream.

A first possibility is that recognizable images contain *category-specific* visual features that are captured by the CNN. As CNNs are trained on millions of exemplars and thousands of categories (e.g., Krihevsky et al., 2012), their convolutional units are both tied to a specific category but inherently visual in nature. These category-specific features could include, for example, different sets of characteristic shape parts that differ between animates and inanimates (e.g., animals tend to have tails, eyes, ears; Levin et al., 2001). There may also be characteristic shape features for small objects (e.g., handles, buttons) and big objects (e.g., extended flat surfaces). This view is broadly in line the idea that the visual feature tunings in occipito-temporal cortex emerge as a byproduct of optimizing for object categorization (e.g., Yamins et al., 2014; Khaligh-Razavi & Kriegeskorte, 2014).

A second possibility is that recognizable images contain additional *generic* visual features that are useful for describing any given object. For example, recognizable images contain strong bounded contours and other visual features that specify their 3D part-structure, whereas texforms do not. Further, a wealth of work has found complex single unit tunings in infero-temporal cortex for both 3D object shape (Hung, Carlson, & Connor, 2012; Yamane, Carlson, Bowman, Wang, & Connor, 2008) and 3D environmental shape (Vaziri, Carlson, Wang, & Connor, 2014). Thus, on an alternative account, it is these kinds of generic visual features, not tied to the category membership of the objects, which account for this differential activity. Certainly, texforms are quite a long way from having any clear three-dimensional structure, so it is a reasonable possibility that the gap can be filled by additional mid-level features that aren’t category-specific per se.

At stake with this distinction is whether the nature of the visual representation in occipito-temporal cortex should be considered more ‘low-level’ or more ‘high-level’ (Peelen & Downing, 2017; Andrews et al., 2015). Interestingly, convolutional neural networks might be able to provide some insight into these questions. For example, if a CNN was trained to perform a simpler task (e.g., a same vs. different image task), then their units would become tuned without any category-specific feedback, but would presumably contain some set of generic visual descriptors. However, perhaps some degree of categorization training (e.g., animate/inanimate, or face/non-face) may be needed to render CNN units complex enough to predict categorical neural responses. Pinpointing the minimal amount of categorization training needed for this structure to emerge remains a rich topic for future work.

## Conclusion

The present work investigated the link between mid-level features and the known animacy and size organizations in occipito-temporal cortex. We found that mid-level feature differences are sufficient to elicit these large-scale organizations along the entire ventral stream. Predictive modeling provided converging support for this result, as both intermediate layers of CNNs and intuitive ratings of curvature predicted the neural pattern similarity elicited by texforms and recognizable images. This work provides new evidence to situate the level of representation in the ventral stream, demonstrating that much of object-selective cortical organization can be explained by relatively primitive mid-level features, without requiring explicit recognition of the objects themselves. Broadly, these data are consistent with the view that the entire ventral stream is predominantly tuned to mid-level visual features, which are spatially organized across the cortex in a way that co-varies with high-level, categorical dimensions.

## Acknowledgements

This work was supported by the Harvard Star Family Challenge grant to T.K and a NIH Shared Instrumentation Grant (S10OD020039) to the Harvard Center for Brain Science.

## Author Contributions

Bria Long and Talia Konkle developed the study idea and B. Long generated and normed stimuli. B. Long collected and analyzed data, using software written by T.K. Chen-Ping Yu executed the CNN training and helped develop the modeling procedure. B. Long, T. Konkle, and C.P. Yu all provided critical revisions to the manuscript.

## Materials and Methods

### Participants

Sixteen healthy observers with normal or corrected-to-normal vision participated in a 2-hour fMRI session (age range 18–35 years; seven females) for Experiment 1 (*N* = 8) and Experiment 2 (*N* = 8). All participants (*N =* 110 across norming and fMRI experiments) were consented using procedures approved by the Institutional Review Board at Harvard University.

### MRI acquisition

Imaging data were collected using a 32-channel phased-array head coil with a 3T Siemens Prisma fMRI Scanner at the Harvard Center for Brain Sciences. High-resolution T1-weighted anatomical scans were acquired using a 3D MPRAGE protocol (176 sagittal slices; FoV = 256 mm; 1x1x1 mm voxel resolution; gap thickness = 0 mm; TR = 2530 ms; TE = 1.69 ms; flip angle = 7 degrees). For functional runs, blood oxygenation level-dependent (BOLD) contrast was obtained using a gradient echo-planar T2* sequence (84 oblique axial slices acquired at a 25° angle off of the anterior commissure-posterior commissure line; FoV = 204 mm; 1.5×1.5×1.5 mm voxel resolution; gap thickness = 0 mm, TR = 2000 ms; TE = 30 ms; flip angle = 80 degrees; multi-band acceleration factor = 3).

### Stimulus Set

The stimulus set consisted of 240 total images, with 120 original images of 30 big animals, 30 big objects, 30 small animals, and 30 small objects, as well as their texform counterparts.

Texforms were created using a texture synthesis algorithm (Freeman & Simoncelli, 2011), in which the original images were placed at peripheral locations with respect to the pooling windows (scaling factor *s*=.5, aspect ratio *AR*=1). These choices were made in order to create relatively spatially coherent stimuli that were not recognizable at the basic-level. See Freeman & Simoncelli (2011) and Long et al. (2016) for further details on the synthesis procedure.

To construct this stimulus set, online norming studies were conducted to on a superset of 240 texforms to ensure they were not recognizable. First, 18 participants guessed the identity of each texform image. Next, six new participants assessed the validity of these guesses. These participants were presented with the original images and all of the texform guesses, and judged whether each guess could be “used to correctly describe” the original image. The proportion of guesses accepted as correct yielded a basic-level identification score for each image. Images in which a rater accepted more than 3/18 responses as correct were removed. Next, 120 texforms and their corresponding originals were selected (30 images per category), with the constraints that the categories did not significantly differ in either aspect ratio or pixel area; all *p* >= .1). On average, these 120 texforms were identified at the basic level <3% of the time. Finally, the overall luminance and contrast levels across all 240 images (120 texforms, 120 originals) equated using the SHINE toolbox (Willenbockel et al., 2010) and the edges of all of images were blurred so that they gradually faded into their backgrounds.

### Texform Classifiability

The classifiability of each texform was calculated using online rating experiments which independently assessed the likelihood that a texform was an animal, an object, big in the world, or small in the world. Specifically, in the first experiment, participants (N=16) were shown a texform and asked: “Here is a scrambled picture of something. Was the original thing an animal?”, and responded with “Yes” or “No”. Similarly, the three other groups of participants (N=16 each) judged whether the texform was a “man-made object”, “big enough to support a human”, and “small enough to hold with one or two hands”.

Animacy and size classifiabilty scores were then calculated from these data by %correct – %incorrect classifications. This serves as a proxy for a d-prime measure and allows for response bias to be factored out from the classification scores. For example, if the texform was generated from an animal original image, this score was calculated as %(yes, was an animal) – %(yes, it’s a man-made object).

Similarly, if the original item was an inanimate object, this score was computed as %(yes, it’s inanimate) – %(yes, it’s an animal). The same procedure was followed for size classifiability: %(yes, it’s big) – %(yes, it’s small) if the original item had a big real-world size; and %(yes, it’s small) – %(yes, it’s big) if the item had a small real-world size. With these measures, the higher the score, the more it was correctly classified as an animal/object and big/small; negative scores indicate systematic misclassifications.

These animacy and size classification scores were summed to obtain a composite classification score. Composite classifiability scores were used to assign the stimuli into 6 groups of 5 images per condition (big animals, big objects, small animals, small objects), from lowest to highest total classifiability. See **Figure S4A** for a visualization of these classification results.

### fMRI Experiment Design

Observers viewed images of big animals, small animals, big objects, and small objects in a standard blocked design while undergoing functional neuroimaging. In the first four runs of the experiment, observers saw texforms; in the second four runs observers saw original images. Observers were not told anything regarding what the texforms were. Unbeknownst to participants, the texform and original runs were yoked, such the original images were shown in the exact same sequence and timing as the texforms. Observers’ task was to pay attention to each item and to press a button when an exact image repeated back to back, which occurred once per block.

Each run had twelve 6s blocks for each condition (big animals, big objects, small animals, small objects), with 10s rest periods interleaved every four blocks. Each block consisted of six images (5 unique images and 1 repeat) each presented for 800ms followed by a 200ms blank. Further, each block contained images from one of the six classifiability levels. Each classifiability level for each condition was shown twice per run. All images were presented in isolation on a uniform gray background.

In Experiment 1, each image subtended 10.36° x 10.36° visual angle centered at fixation. In Experiment 2, each block of images could appear either above or below fixation (6.92° x 6.92° degrees of visual angle, bottom edge .86° degrees from center). These positions were counterbalanced across blocks such that, for each level of classifiability and condition, one block was presented in the upper visual field and the other block was presented in the lower visual field. Participants were instructed that maintaining fixation was more important than task performance, and fixation was monitored online using an EyeLink 1000 eye-tracker. Participants were calibrated to the eye-tracker at the beginning of the experiment and were recalibrated every 2-3 runs as needed.

### Preference Map Correlations

The spatial distribution and strength of response preferences in visually-active voxels along the ventral stream using a preference-map analysis (see Konkle and Oliva, 2012b; Konkle & Caramazza, 2013). Active voxels were selected from a contrast of all conditions > rest with *t* > 2 in each subject separately for both the texforms runs and the original runs, and then these active texform and original masks were intersected. For the animacy organization, for each voxel, the betas for both big and small animals were averaged and subtracted from the betas for both big and small objects; this beta difference map was displayed on the cortical surface within an active voxel mask. For the size organization, for each voxel, the beta for big objects was subtracted from the beta for small objects, masked, and displayed on the cortical surface.

To compare animacy and real-world size preference maps elicited by texform and original images, we used a map-correlation procedure, similar to Srihasam et al., 2014. First, we defined the set of voxels over which to compare the original and texform maps. This set was defined in each subject as the set of all active voxels (all conditions > rest with *t* > 2), falling within the occipital and temporal lobes and their gray matter mask, excluding voxels within functionally-defined early visual areas V1-V3. Within these voxels, the animacy map-correlation was computed as the correlation between the beta-difference scores for the texform animacy organization and original animacy organization. For the size organization, the same analysis was performed but using big–small object beta differences. These map correlations were separately computed separately for each subject. For statistical comparisons, single-subject r-values were fischer-z transformed and entered into a t-test.

A split-half analysis was used to estimate the noise ceiling on the map-correlations, for both animacy and real-world size organizations. Specifically, in each subject, texform preference maps were correlated between odd and even runs, yielding a texform map split-half correlation. The same analysis was repeated for originals. If any of these odd-even correlations was less than zero (i.e., a negative correlation), we substituted this value with zero; this occurred in one subject for the object size comparison. These reliabilities were then corrected using the Spearman Brown prophecy formula (*N*\*observed reliability / 1 +(*N*-1)*observed reliability, where *N* = 2 as we divided the data in half). The noise ceiling in each participant was computed taking the square root of the product of these corrected reliabilities; a paired t-test was used to compare subject’s actual correlation value with their individual noise ceilings.

### Posterior to Anterior Analyses

In each participant, anatomical sections were defined along a posterior to anterior gradient within occipito-temporal cortex, by dividing it into five quantiles using the TAL-Y coordinates of visually active voxels (taken from each participant’s GLM data). Next, within each section, we identified the voxels that had the same animacy preference between originals and texforms (e.g., a voxel preferred both recognizable animals and texform animals). Within these comparable voxels, we obtained a measure of the strength of the animacy preferences for either objects or texforms, computed as the absolute value of animals-object betas for each voxel, averaged across voxels. These estimates were computed separately for originals and texforms, in each section, in each participant.

To assess whether there was a difference in the overall strength of the original animacy preferences vs. the texform animacy preferences along the posterior-to-anterior gradient, we computed difference scores for each anatomical section for each participant. We then performed a simple rank correlation between ascending anatomical sections (i.e., 1,2,3,4,5) and these difference scores (orig – texforms) in each subject. A rank correlation metric was used to assess whether originals generate greater animacy preferences along this posterior to anterior gradient, without assuming a meaningful relationship with the TAL-Y coordinates of the anatomical sections. Finally, we asked whether these rank correlations were above zero at the group level by performing a one-sample t-test over subjects. The same procedure was repeated for the object size distinction. For visualization purposes, in **Figure 3A-C**, we also defined group-level anatomical sections based on the Group GLM activations, with the scatter plots showing voxel response preferences for animacy and size dimensions based on the group GLM beta fits.

### Conjunction Analysis

Conjunction voxels were defined as those that elicited the same category preference (e.g., animals) regardless of the location of the image in the visual field (i.e., upper visual field, lower visual field) in response to recognizable images (i.e., originals). Conjunction voxels were defined separately within each subject. To calculate the portion of retained voxels, we divided the number of voxels in this conjunction mask by the total number of visually active voxels in OTC in each subject. We then performed map correlations in each subject within these conjunction voxels.

### Representational Similarity Analysis

Multi-voxel patterns were extracted for each of the four main conditions (animals/objects x big/small) at each level of classifiability (levels 1-6), yelling 24 conditions. To reduce noise, multi-voxel patterns were only extracted in voxels where recognizable images yielded a split-half reliability value above zero. Next, the pairwise similarity among neural patterns was computed using a correlation metric r, converted to correlation distance by taking 1-r, and arranged into square representational dissimilarity matrices (RDMs) for visualization purposes. RDMs were computed separately for texforms and original images, for each participant. Comparisons between RDMs were computed by taking the correlation of the elements in the lower-diagonal matrix of the RDMs, and all statistical tests were conducted on fisher-z transformed r-values.

### Predictive Modeling Approach

To compare how well different models (i.e., image statistics models, CNN features, and behavioral ratings) could predict the neural RDMs, we used the predictive modeling procedures used by Jozwik et al. (2016) as implemented in the RSA toolbox (Nili et al., 2014). This method finds the weighted combination of feature dissimilarity matrixes (e.g., RDMs) that best predict a subset of the neural data using non-negative least squares regression with leave-one-condition-out cross-validation (NNLS; Lawson & Hanson, 1974). A neural dissimilarity vector (276x1) was constructed iteratively over cross-validation loops. The goodness-of-prediction was assessed by correlating between the actual neural RDM from each individual subject with the cross-validated model RDM. The noise ceiling was computed using the RSA toolbox (Nili et al., 2014), reflecting the degree to which an individual subject’s RDM could predict the group’s RDM. See Supplemental Information: Methods and Materials for descriptions how features were extracted from different models for use in this predictive modeling procedure.

### Open-Access Data & Code

All stimuli, pre-processed data, and main analysis code for this paper are available at the Open Science Repository for this project, https://osf.io/69pbd/, which is also linked to a GitHub codebase for generating texforms. Raw fMRI data is available on request.

## Supplemental Information

### Supplemental Methods

#### Data preprocessing

Functional data were analyzed using Brain Voyager QX software and MATLAB. Preprocessing included slice scan-time correction, 3D motion correction, linear trend removal, temporal high-pass filtering (0.01 Hz cutoff), spatial smoothing (4 mm FWHM kernel), and transformation into Talairach (TAL) coordinates. Two subjects had one run in which they moved more than 0.5 mm within 2 seconds (1 TR) and these runs were discarded from analysis. The cortical surface of each subject from the high-resolution T1-weighted anatomical scan, acquired with a 3D MPRAGE protocol. To do so, we used the default segmentation procedures in FreeSurfer. Surfaces were then imported into Brain Voyager and inflated using BV surface module. Gray matter masks were defined in the volume and were constructed based on the Freesurfer segmentations.

General linear models (GLMs) were computed at the single subject level for texforms and original runs separately, both for the four main conditions (big animals, big objects, small animals, and small objects) as well as separately for the full set of nested conditions (each category x each classifiability level, 24 conditions total). GLMs included square-wave regressors for each condition’s presentation times, convolved with a gamma function to approximate the hemodynamic response. In Experiment 2, GLMs were fit eight main conditions of interest: each combination of category (big animals, big objects, small animals, and small objects) and visual field presentation (upper, lower) separately for texforms and originals.

#### Retinotopy Protocol

Additionally, participants completed a retinotopy protocol in order to define early visual areas V1-V3. Observers viewed bands of flickering checkerboards in a blocked design. The conditions included vertical meridian bands (~22° × 1.7°), horizontal meridian bands (~22° × 1.7°), iso-eccentricity bands covered by a central ring (radius ~1.2° to 2.4°), a peripheral ring (radius ~5.7° to 9.3°), and an extra wide peripheral ring (inner radius ~9.3°, filling the extent of the screen). In Experiment 2, the vertical and horizontal meridian bands were replaced with wedges. The apex of each wedge was at fixation and the base extended to ~22° in the periphery, and the checkerboard patterns flickered at 6 Hz. Each block was 6 seconds, within which the checkerboard cycled at 8 Hz between states of black-and-white, randomly colored, white-and-black, and random colored. In each 4.4-min run (142 volumes), the 5 visual field band conditions and 1 fixation condition were repeated 7 times with their order randomly permuted within each repetition. Each run started and ended with a 6 s fixation period. Participants’ task was to maintain fixation, and press a button every time the fixation dot turned red, which happened once per block. Using data from this retinotopy protocol, early visual regions (V1-V3) were defined by hand on inflated brain guided by the contrast of horizontal vs. vertical meridians (see Cohen et al., 2014).

### Predictive Modeling Feature Spaces

#### Gabor & Gist Models

Gabor features were extracted in an 8 × 8 grid over the original, recognizable images (440 × 440 pixels) at three different 3 scales, with 8,6, and 4 oriented Gist per scale, respectively (Oliva & Torralba, 2006). GIST model features were extracted by taking the first 20 principle components of this Gabor feature matrix. In both cases, these features were then averaged across the five images in each nested classifiability group presented during the fMRI experiment. The squared Euclidean distance along each feature was used to construct feature RDMs for use in the predictive modeling procedure, and all dissimilarities were scaled between 0 and 1, yielding a 276 × 896 feature vector for Gabor features and a 276 × 20 feature vector for Gist features.

#### Texture Synthesis Model (Freeman & Simoncelli, 2011)

The texture synthesis model has 10 feature classes (corresponding to pixel statistics, weighted marginal statistics, simple cell responses, complex cell responses, cross-position correlations (i.e., autocorrelation) within scales computed separately for simple and complex cells, cross-orientation correlations computed separately for simple and complex cells, and cross-scale correlations computed separately for simple and complex cells). Features were included if they (1) had any variance across the images (SD > 0) and (2) were calculated within pooling windows tiling the depicted item. The values for each feature were then z-scored across the 120 images, and then averaged over the five images in each classifiability group. Each feature was converted to an RDM using squared Euclidean distance, and all dissimilarities were scaled between 0 and 1, generating a 276 × 20,914 feature matrix.

#### Behavioral Ratings–Animacy/Size

For texforms, feature RDMs were constructed based on the behavioral animacy and size classifiability scores. Note these are the same scores used to group the texforms into the nested design. These experiments yielded a vector corresponding to participants ability to classify each texform as an animal (range: 0-1, where 1 = always classified as an animal, and 0 = never classified as an animal) and their ability to classify each texform according to their size in the real world (range: 0-1, where 1=always classified as big in the real-world, and 0 = never classified as big in the real world). These scores were averaged according to the 24 nested conditions presented during the experiment, yielding a 24 × 1 vector for animacy and a 24 × 1 vector for size for texforms and for originals. We then took the squared Euclidean distance of each 1-dimensional feature vector and all dissimilarities were scaled between 0 and 1; the final feature matrix was a 276 × 2 feature matrix.

For original images, feature RDMs were constructed using their actual animacy/size in real-world. These yielded a vector corresponding to the actual animacy of the recognizable image (1=animate, and 0=inanimate), and a vector corresponding the actual size of the object in the real world (1=big in the real-world, and 0=small in the real world). We then took the squared Euclidean distance of each 1-dimensional feature vector and all dissimilarities were scaled between 0 and 1; the final feature matrix was a 276 × 2 feature matrix.

#### Behavioral Ratings–Curvature

Behavioral ratings on Amazon Mechanical Turk were obtained to assess the perceived curvature of both the texforms and the originals; 30 participants rated the curvature of the 120 texforms, and another 30 participants rated the curvature of their 120 corresponding original images. Participants were asked, “How boxy or curvy is the thing depicted in this image?” and asked to respond using a 1-5 scale (1: Very curvy, 2: Mostly curvy, 3: Equally boxy and curvy, 4: Mostly boxy, 5: Very boxy). These ratings were averaged across participants, and then averaged across the five images in each classification group. This yielded two 24 × 1 vectors corresponding to the average perceived curvature of each group of texforms and of each group of original images. The squared Euclidean distance of all pairwise comparisons of these conditions was computed separately for texforms and originals, yielding two 276 × 1 feature vectors for curvature for modeling responses to texforms and originals; all dissimilarities were scaled between 0 and 1. See **Figure S4B** for a visualization of this data and a comparison of the curvature ratings between texforms and recognizable images.

#### CNN Features (Texforms & Originals)

The AlexNet architecture (Krizhevsky et al., 2012) as was trained using the conventional image classification task using the ImageNet dataset (Russakovsky et al., 2015). The standard AlexNet training regime was adopted using a public code package (https://github.com/soumith/imagenet-multiGPU.torch) that was optimized for multi-threaded CNN training in Torch7. Specifically, stochastic gradient descent (SGD) optimization was used with 0.9 momentum, an initial learning rate of 0.02, and weight decay of 0.0005. Both the learning rate and the weight decay follow a pre-defined decreasing schedule (see train.lua from the code package) using a mini-batch size of 128, with 10,000 mini-batches per epoch over a total of 55 training epochs. Standard data augmentation such as random horizontal flips and random 224-by-224 crops were performed during training.

Using the fully-trained network, CNN features were extracted from each unit in the CNN from both original and texform image sets. Specifically, for each image and each convolutional filter, we computed the summed activation map of the filter (an m-by-m map where m is the output size of the convolutional layer), accounting for border effects by setting to zero all values in the activation map within 10% of the four edges. This procedure yielded five feature matrices for the original images of 120-by-64, 120-by-192, 120-by-384, 120-by-256, and 120-by-256, corresponding to each of the five convolutional layers, and another set corresponding to the texform images. For the two fully-connected layers (layer 6 & 7), the activation level to each image was direct computed (no global summation required), resulting in two feature matrices of 120-by-4096, corresponding to layer 6 and 7, and another set for texform images.

Each feature matrix was normalized by dividing the rows with its L2-norm, and the rows were averaged over the five images in each classifiability group. Finally, for each feature (column) of each feature matrix, we computed the pairwise squared Euclidean distance between all of the 24 conditions, yielding five 276-by-m representational dissimilarity matrices, where m is the number of convolutional filters for the corresponding layer. All dissimilarities were then scaled between 0 and 1. These RDMs were used to perform feature modeling of the individual CNN layers.

---

**Supplemental Figure 1. Related to Figure 2.**
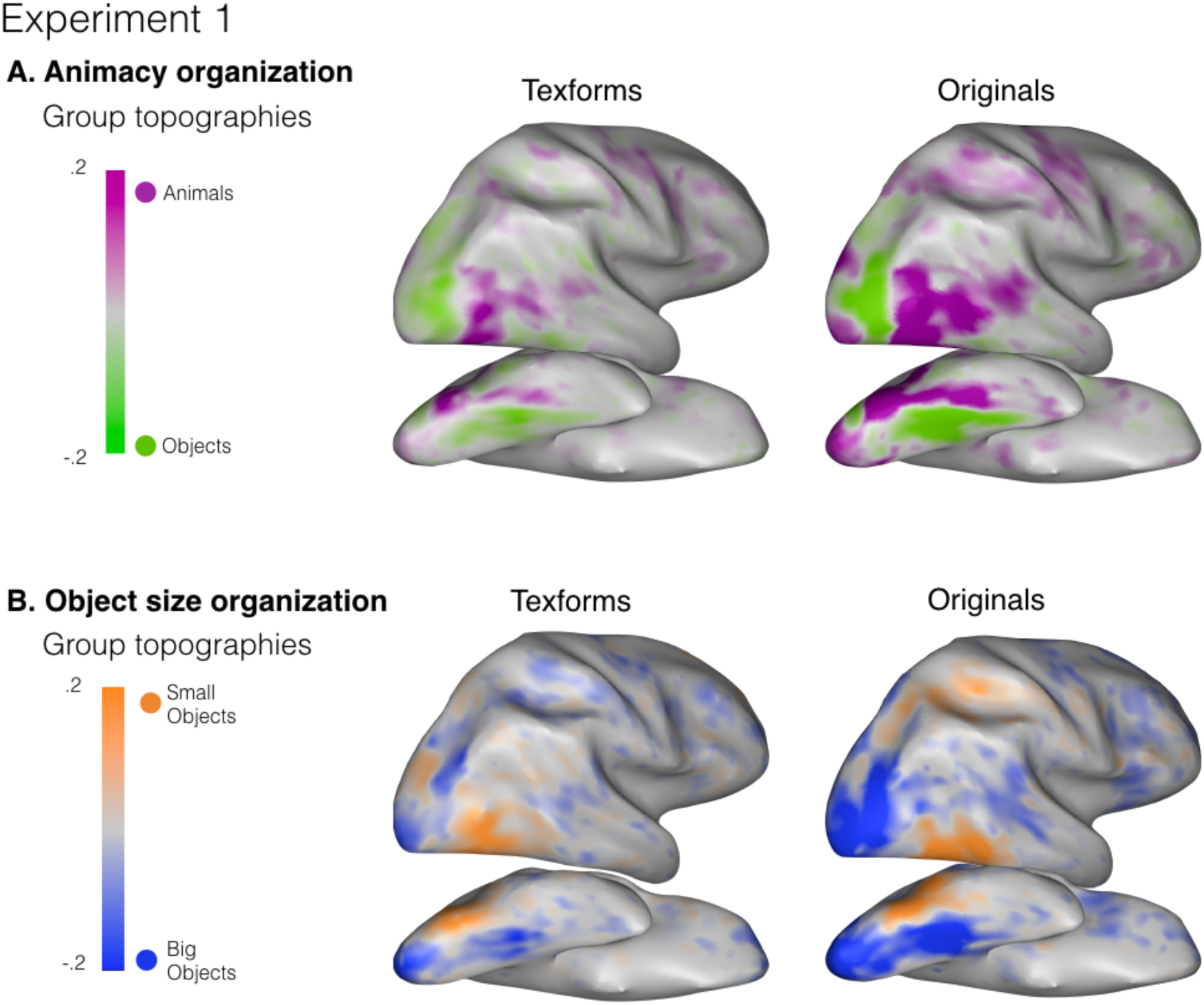
Group topographies for both the animacy (A) and object size (B) contrasts are shown. Voxels shown include any voxel where any subject had an active voxel (all >rest, *t* > 2); group topographies are projected onto an individual’s anatomical brain.

**Supplemental Figure 2. Related to Figure 4.**
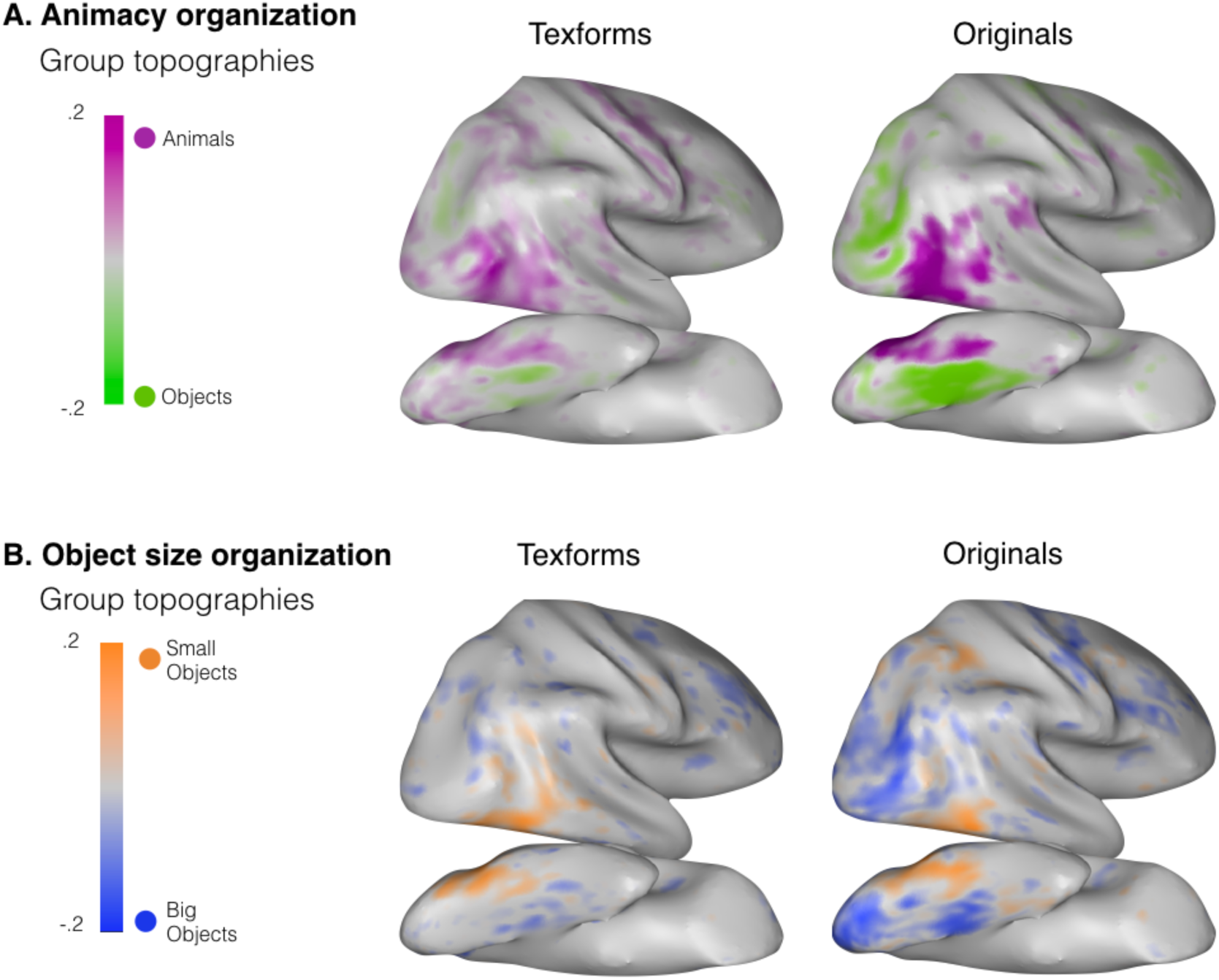
Group conjunction topographies for both the animacy (A) and object size (B) contrasts are shown. These were computed topographies were computed in the same manner for original subjects but using the group GLM. Voxels shown include any voxel where any subject had an active voxel (all>rest, *t*>2); group topographies are projected onto an individual’s anatomical brain.

**Supplemental Figure 3. Related to Figure 5.**
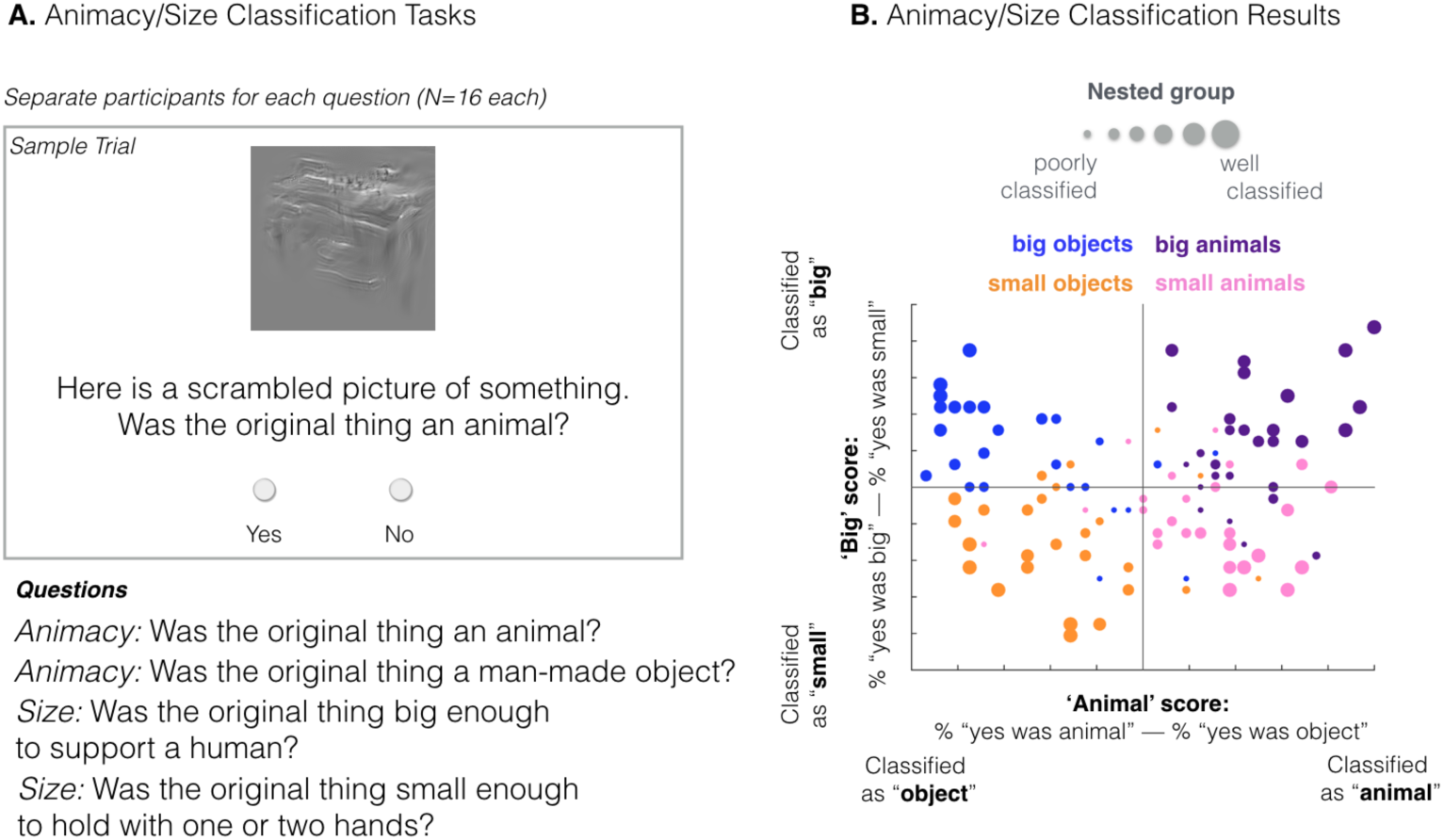
(A) Schematic of the animacy/size classification tasks. (B)The classifiability each image plotted for animacy (x-axis) and real-world size (y-axis); the dot size corresponds to the nested classifiability group in which this image was included; larger dots represent groups of images that were better classified by their animacy and real-world size.

**Supplemental Figure 4. Related to Figure 6.**
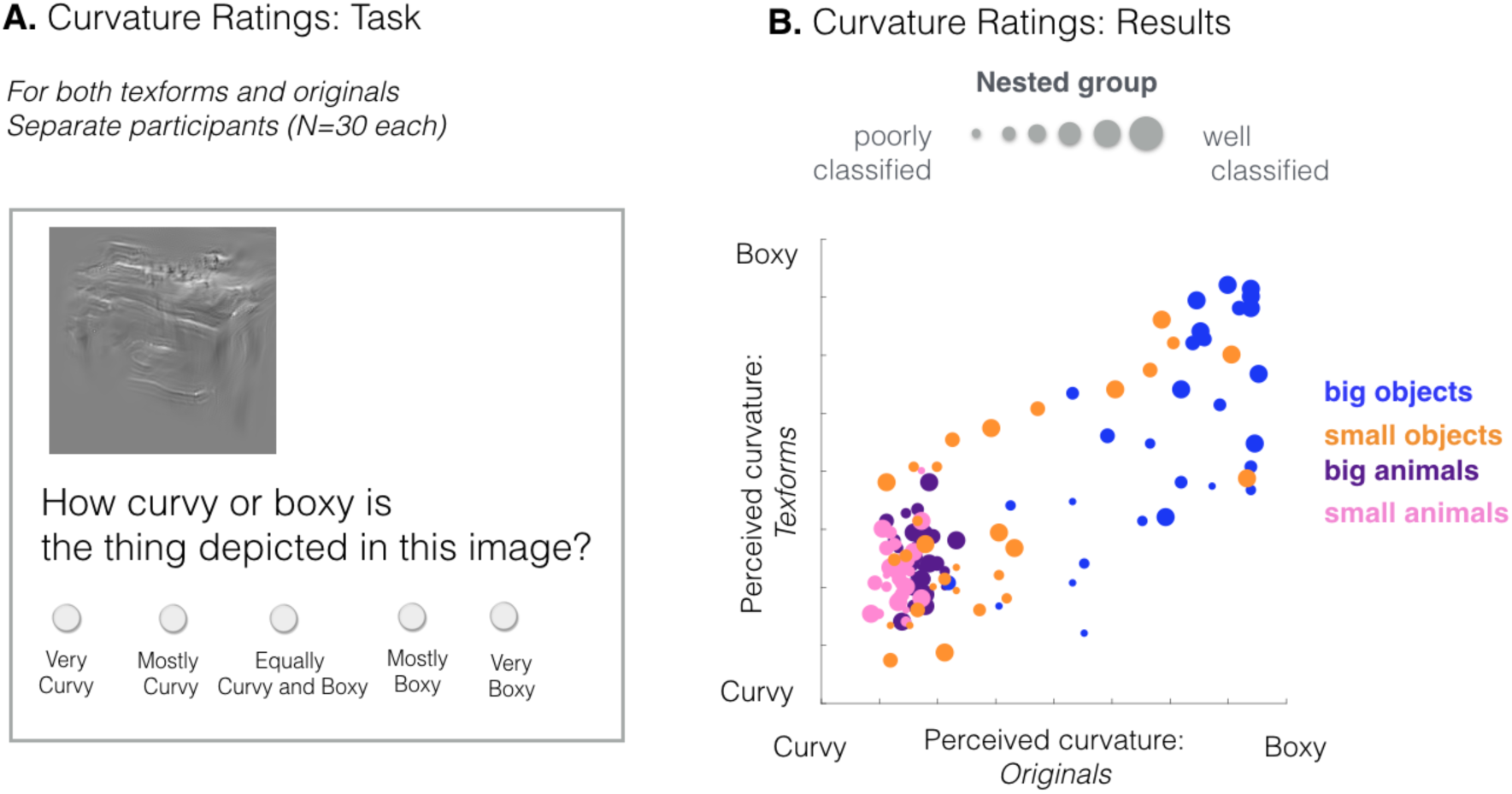
(A) Schematic of the perceived curvature task. (B). The perceived curvature of the texforms (y-axis) is plotted as a function of the perceived curvature of the recognizable, original images (x-axis); larger dots represent groups of images that were better classified by their animacy and real-world size.

**Supplemental Figure 5. Related to Figure 6.**
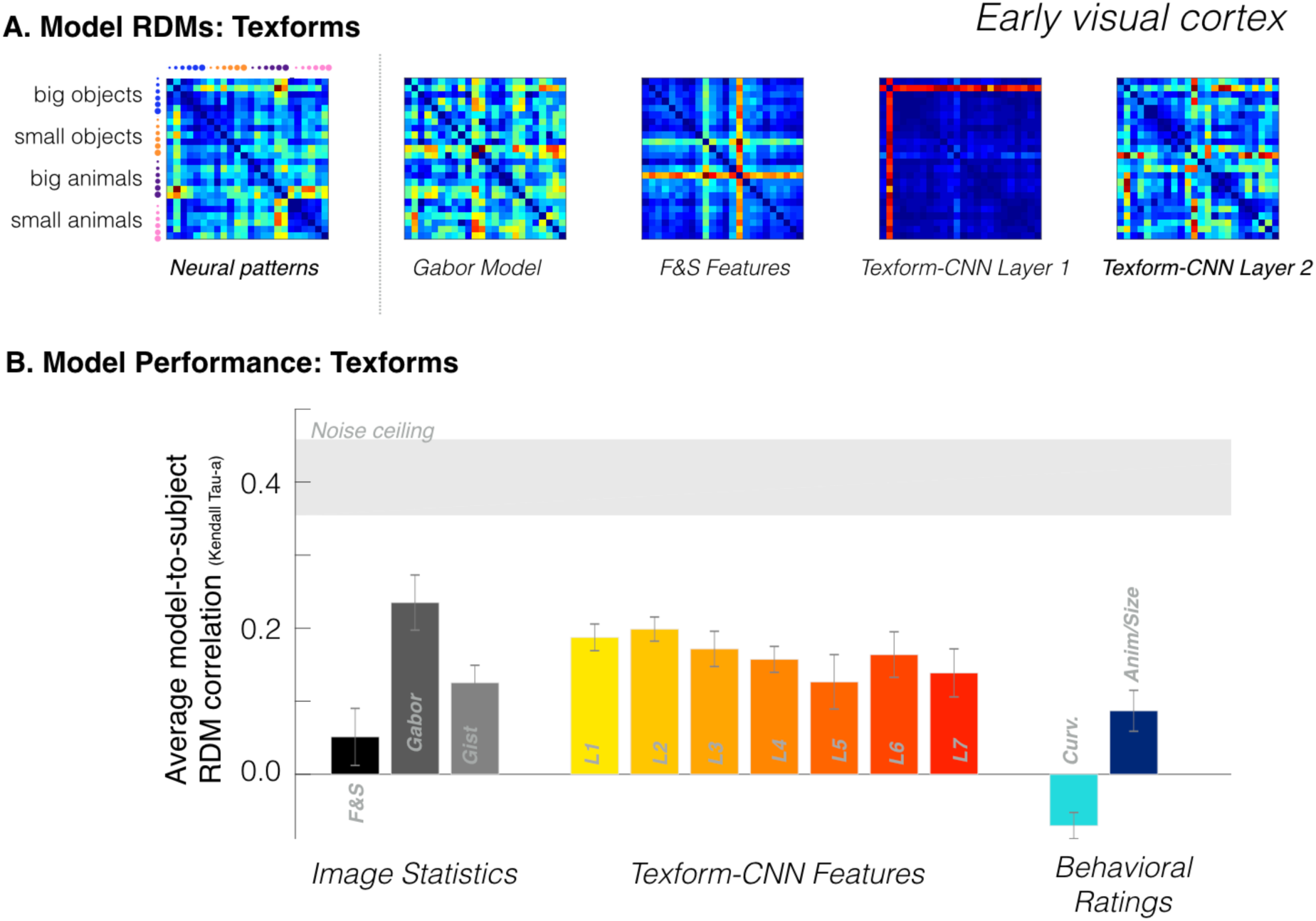
(A) Neural patterns in response to texforms in *early visual cortex* and predicted neural dissimilarities for selected models obtained through the same cross-validation procedure. (B) Average predicted model correlation (kendall tau-a) with individual subjects’ neural patterns in EVC. Data is plotted with respect to the noise ceiling, shown in light gray.

## References

Andrews, T. J., Watson, D. M., Rice, G. E., & Hartley, T. (2015). Low-level properties of natural images predict topographic patterns of neural response in the ventral visual pathway. Journal of Vision, 15(7), 3.

Andrews, T. J., Clarke, A., Pell, P., & Hartley, T. (2010). Selectivity for low-level features of objects in the human ventral stream. Neuroimage, 49(1), 703–711.

Baldassi, C., Alemi-Neissi, A., Pagan, M., DiCarlo, J. J., Zecchina, R., & Zoccolan, D. (2013). Shape similarity, better than semantic membership, accounts for the structure of visual object representations in a population of monkey inferotemporal neurons. PLoS Computational Biology, 9(8), e1003167.

Bi, Y., Wang, X., & Caramazza, A. (2016). Object Domain and Modality in the Ventral Visual Pathway. Trends in Cognitive Sciences, 20(4), 282–290.

Biederman, I. (1987). Recognition by components: A theory of human image understanding. Psychological Review, 94(2), 115–117.

Bracci, S., & Op de Beeck, H. (2016). Dissociations and Associations between Shape and Category Representations in the Two Visual Pathways. Journal of Neuroscience, 36(2), 432–444.

Brewer, A. A., Liu, J., Wade, A. R., & Wandell, B. A. (2005). Visual field maps and stimulus selectivity in human ventral occipital cortex. Nature Nemuroscience, 8(8), 1102–1109.

Bryan, P., Julian, J., & Epstein, R. (2016). Rectilinear edge selectivity is insufficient to explain the category selectivity of the parahippocampal place area. Frontiers in Human Neuroscience. Retrieved from

Castelli, F., Happé, F., Frith, U., & Frith, C. (2000). Movement and mind: a functional imaging study of perception and interpretation of complex intentional movement patterns. NeuroImage, 12(3), 314–25.

Chao, L. L., Haxby, J. V, & Martin, A. (1999). Attribute-based neural substrates in temporal cortex for perceiving and knowing about objects. Nature Neuroscience, 2(10), 913–919.

Cohen, L., Dehaene, S., Naccache, L., Lehéricy, S., Dehaene-Lambertz, G., Hénaff, M.-A., & Michel, F. (2000). The visual word form area: spatial and temporal characterization of an initial stage of reading in normal subjects and posterior split-brain patients. Brain, 123(2), 291–307.

Coggan, D. D., Liu, W., Baker, D. H., & Andrews, T. J. (2016). Category-selective patterns of neural response in the ventral visual pathway in the absence of categorical information. NeuroImage, 135, 107–114.

Güçlü, U., & van Gerven, M. A. J. (2015). Deep Neural Networks Reveal a Gradient in the Complexity of Neural Representations across the Ventral Stream. Journal of Neuroscience, 35(27), 10005–10014.

DiCarlo, J. J., & Cox, D. D. (2007). Untangling invariant object recognition. Trends in Cognitive Sciences, 11(8), 333–341.

Downing, P. E., Jiang, Y., Shuman, M., & Kanwisher, N. (2001). A cortical area selective for visual processing of the human body. Science, 293(5539), 2470–2473.

Epstein, R., & Kanwisher, N. (1998). A cortical representation of the local visual environment. Nature, 392(6676), 598–601.

Felleman, D. J., & Van Essen, D. C. (1991). Distributed hierachical processing in the primate cerebral cortex. Cerebral Cortex, 1(1), 1–47.

Freeman, J., & Simoncelli, E. P. (2011). Metamers of the ventral stream. Nature Neuroscience, 14(9), 1195–1201.

Freeman, J., Ziemba, C. M., Heeger, D. J., Simoncelli, E. P., & Movshon, J. A. (2013). A functional and perceptual signature of the second visual area in primates. Nature Neuroscience, 16(7), 974–81.

Goodale, M. A., & Milner, A. D. (1992). Separate visual pathways for perception and action. Trends in Neurosciences, 15(1), 20–25.

Golomb, J. D., & Kanwisher, N. (2012). Higher level visual cortex represents retinotopic, not spatiotopic, object location. Cerebral Cortex, 22(12), 2794–2810. https://doi.org/10.1093/cercor/bhr357

Grill-Spector, K., & Weiner, K. S. (2014). The functional architecture of the ventral temporal cortex and its role in categorization. Nature Reviews Neuroscience Neuroscience, 15(8), 536–548.

Hasson, U., Levy, I., Behrmann, M., Hendler, T., & Malach, R. (2002). Eccentricity bias as an organizing principle for human high-order object areas. Neuron, 34(3), 479–490.

Haxby, J. V., Gobbini, M. I., Furey, M. L., Ishai, A., Schouten, J. L., & Pietrini, P. (2001). Distributed and Overlapping Representations of Faces and Objects in Ventral Temporal Cortex. Science, 293(5539), 2425– 2425.

He, C., Peelen, M. V., Han, Z., Lin, N., Caramazza, A., & Bi, Y. (2013). Selectivity for large nonmanipulable objects in scene-selective visual cortex does not require visual experience. NeuroImage, 79, 1–9.

Hong, H., Yamins, D. L. K., Majaj, N. J., & DiCarlo, J. J. (2016). Explicit information for category-orthogonal object properties increases along the ventral stream. Nature Neuroscience, 19(4), 613–622.

Hung, C.-C., Carlson, E. T., & Connor, C. E. (2012). Medial Axis Shape Coding in Macaque Inferotemporal Cortex. Neuron, 74(6), 1099–1113. https://doi.org/10.1016/j.neuron.2012.04.029

Jozwik, K., Kriegeskorte, N., & Mur, M. (2016). Visual features as stepping stones toward semantics: Explaining object similarity in IT and perception with non-negative least squares. Neuropsychologia.

Julian, J. B., Ryan, J., & Epstein, R. A. (2016). Coding of Object Size and Object Category in Human Visual Cortex. Cerebral Cortex, 29, 1–15.

Kaiser, D., Azzalini, D. C., & Peelen, M. V. (2016). Shape-independent object category responses revealed by MEG and fMRI decoding. Journal of Neurophysiology, 115(4), 2246–2250.

Kanwisher, N., McDermott, J., & Chun, M. M. (1997). The fusiform face area: a module in human extrastriate cortex specialized for face perception. Journal of Neuroscience, 17(11), 4302–4311.

Khaligh-Razavi, S. M., & Kriegeskorte, N. (2014). Deep Supervised, but Not Unsupervised, Models May Explain IT Cortical Representation. PLoS Computational Biology, 10(11), e1003915.

Konkle, T., & Caramazza, A. (2013). Tripartite Organization of the Ventral Stream by Animacy and Object Size. Journal of Neuroscience, 33(25), 10235–10242.

Konkle, T., & Caramazza, A. (2016). The Large-Scale Organization of Object-Responsive Cortex Is Reflected in Resting-State Network Architecture. Cerebral Cortex, 31(28), 1–13.

Konkle, T., & Oliva, A. (2012). A Real-World Size Organization of Object Responses in Occipitotemporal Cortex. Neuron, 74(6), 1114–1124.

Kourtzi, Z., & Connor, C. E. (2010). Neural Representations for Object Perception: Structure, Category, and Adaptive Coding. Annual Review of Neuroscience, 34(1), 45–67.

Kravitz, D. J., Saleem, K. S., Baker, C. I., Ungerleider, L. G., & Mishkin, M. (2013). The ventral visual pathway: an expanded neural framework for the processing of object quality. Trends in Cognitive Sciences, 17(1), 26–49.

Krizhevsky, A., Sulskever, Ii., & Hinton, G. E. (2012). ImageNet Classification with Deep Convolutional Neural Networks. NIPS, 1–9.

Larsson, J., & Heeger, D. J. (2006). Two retinotopic visual areas in human lateral occipital cortex. The Journal of Neuroscience: The Official Journal of the Society for Neuroscience, 26(51), 13128–42.

Lerner, Y., Harel, M., & Malach, R. (2004). Rapid completion effects in human high-order visual areas. Neuroimage, 21(2), 516–526.

Levin, D. T., Takarae, Y., Miner, a G., & Keil, F. (2001). Efficient visual search by category: specifying the features that mark the difference between artifacts and animals in preattentive vision. Perception & Psychophysics, 63(4), 676–697.

Levy, I., Hasson, U., Avidan, G., Hendler, T., & Malach, R. (2001). Center-periphery organization of human object Lerner, Y., Harel, M., & Malach, R. (2004). Rapid completion effects in human high-order visual areas. Neuroimage, 21(2), 516–526.

Long, B., Konkle, T., Cohen, M. A., & Alvarez, G. A. (2016). Mid-level perceptual features distinguish objects of different real-world sizes. Journal of Experimental Psychology: General, 145(1), 95–109.

Martin, A. (2007). The representation of object concepts in the brain. Annu. Rev. Psychol., 58, 25–45.

Mishkin, M., Ungerleider, L. G., & Macko, K. A. (1983). Object vision and spatial vision: two cortical pathways. Trends in Neurosciences, 6, 414–417.

Murty, N. A. R., & Pramod, R. T. (2016). To What Extent Does Global Shape Influence Category Representation in the Brain? Journal of Neuroscience, 36(15), 4149–4151.

Nasr, S., Echavarria, C. E., & Tootell, R. B. H. (2014). Thinking Outside the Box: Rectilinear Shapes Selectively Activate Scene-Selective Cortex. Journal of Neuroscience, 34(20).

Nili, H., Wingfield, C., Walther, A., Su, L., Marslen-Wilson, W., & Kriegeskorte, N. (2014). A Toolbox for Representational Similarity Analysis. PLoS Computational Biology, 10(4), e1003553.

Okazawa, G., Tajima, S., & Komatsu, H. (2015). Image statistics underlying natural texture selectivity of neurons in macaque V4. Proceedings of the National Academy of Sciences of the United States of America, 112(4), E351–60.

Oliva, A., & Torralba, A. (2006). Building the gist of a scene: The role of global image features in recognition. Progress in Brain Research, 155, 23–36.

Op de Beeck, H. P., Torfs, K., & Wagemans, J. (2008). Perceived shape similarity among unfamiliar objects and the organization of the human object vision pathway. Journal of Neuroscience, 28(40), 10111–10123.

Peelen, M. V, & Downing, P. E. (2017). Category selectivity in human visual cortex: Beyond visual object recognition. Neuropsychologia.

Peelen, M. V, He, C., Han, Z., Caramazza, A., & Bi, Y. (2014). Nonvisual and visual object shape representations in occipitotemporal cortex: evidence from congenitally blind and sighted adults. Journal of Neuroscience, 34(1), 163–170.

Proklova, D., Kaiser, D., & Peelen, M. V. (2016). Disentangling Representations of Object Shape and Object Category in Human Visual Cortex: The Animate–Inanimate Distinction. Journal of Cognitive Neuroscience, 28(5), 680–692.

Lambon Ralph, M. A.., Jefferies, E., Patterson, K., & Rogers, T. T. (2016). The neural and computational bases of semantic cognition. Nature Reviews Neuroscience.

Lehky, S. R., & Tanaka, K. (2016). Neural representation for object recognition in inferotemporal cortex. Current Opinion in Neurobiology, 37, 23–35. https://doi.org/10.1016/j.conb.2015.12.001

Pasupathy, A., & Connor, C. E. (2001). Shape representation in area V4: position-specific tuning for boundary conformation. Journal of Neurophysiology, 86(5), 2505–2519.

Pasupathy, A., & Connor, C. E. (2002). Population coding of shape in area V4. Nature Neuroscience, 5(12), 1332–1338. https://doi.org/10.1038/nn972

Ponce, C. R., Hartmann, T. S., & Livingstone, M. S. (2017). End-stopping predicts curvature tuning along the ventral stream. Journal of Neuroscience, 37(3), 648–659.

Rajimehr, R., Devaney, K. J., Bilenko, N. Y., Young, J. C., & Tootell, R. B. H. (2011). The “Parahippocampal Place Area” Responds Preferentially to High Spatial Frequencies in Humans and Monkeys. PLOS Biology, 9(4),

Rajimehr, R., Bilenko, N. Y., Vanduffel, W., & Tootell, R. B. H. (2014). Retinotopy versus face selectivity in macaque visual cortex. Journal of Cognitive Neuroscience, 26(12), 2691–700.

Ritchie, J. B., Bracci, S., & de Beeck, H. O. (2017). Avoiding illusory effects in representational similarity analysis: What (not) to do with the diagonal. Elsevier.

Rust, N. C., & Dicarlo, J. J. (2010). Selectivity and tolerance (“invariance”) both increase as visual information propagates from cortical area V4 to IT. The Journal of Neuroscience: The Official Journal of the Society for Neuroscience, 30(39), 12978–12995.

Schwarzlose, R. F., Swisher, J. D., Dang, S., & Kanwisher, N. (2008). The distribution of category and location information across object-selective regions in human visual cortex. Proceedings of the National Academy of Sciences of the United States of America, 105(11), 4447–52.

Srihasam, K., Vincent, J. L., & Livingstone, M. S. (2014). Novel domain formation reveals proto-architecture in inferotemporal cortex. Nature Neuroscience, 17(12), 1776–1783.

Striem-Amit, E., & Amedi, A. (2014). Visual cortex extrastriate body-selective area activation in congenitally blind people “seeing” by using sounds. Current Biology, 24(6), 687–692.

Tanaka, K. (2003). Columns for complex visual object features in the inferotemporal cortex: clustering of cells with similar but slightly different stimulus selectivities. Cerebral Cortex, 13(1), 90–99. https://doi.org/10.1093/cercor/13.1.90

van den Hurk, J., Van Baelen, M., & de Beeck, H. P. O. (2017). Development of visual category selectivity in ventral visual cortex does not require visual experience. Proceedings of the National Academy of Sciences, 201612862.

Vaziri, S., Carlson, E. T., Wang, Z., & Connor, C. E. (2014). A channel for 3D environmental shape in anterior inferotemporal cortex. Neuron, 84(1), 55–62.

Weisberg, J., Van Turennout, M., & Martin, A. (2006). A neural system for learning about object function. Cerebral Cortex, 17(3), 513–521.

Wheatley, T., Milleville, S. C., & Martin, A. (2007). Understanding Animate Agents. Psychological Science, 18(6), 469–474.

Yamane, Y., Carlson, E., Bowman, K., & Wang, Z. (2008). A neural code for three-dimensional object shape in macaque inferotemporal cortex. Nature. Retrieved from http://www.nature.com/neuro/journal/v11/n11/abs/nn.2202.html

Yamins, D. L. K., Hong, H., Cadieu, C. F., Solomon, E. A., Seibert, D., & DiCarlo, J. J. (2014). Performance-optimized hierarchical models predict neural responses in higher visual cortex. Proceedings of the National Academy of Sciences of the United States of America, 111(23).

Yau, J. M., Pasupathy, A., Brincat, S. L., & Connor, C. E. (2012). Curvature processing dynamics in macaque area V4. Cerebral Cortex, 23(1), 198–209.

